# Consequences of PDGFRα^+^ fibroblast reduction in adult murine hearts

**DOI:** 10.1101/2021.05.07.443103

**Authors:** Jill T. Kuwabara, Sumit Bhutada, Vikram Shettigar, Greg S. Gojanovich, Lydia P. DeAngelo, Jack R. Heckl, Julia R. Jahansooz, Dillon K. Tacdol, Mark T. Ziolo, Suneel S. Apte, Michelle D. Tallquist

## Abstract

Fibroblasts produce the majority of collagen in the heart and are thought to regulate extracellular matrix (ECM) turnover. Although fibrosis accompanies many cardiac pathologies and is generally deleterious, the role of fibroblasts in maintaining the basal ECM network and in fibrosis *in vivo* is poorly understood. We genetically ablated fibroblasts in mice to evaluate the impact on homeostasis of adult ECM and cardiac function after injury. Fibroblast-ablated mice demonstrated a 60-80% reduction in cardiac fibroblasts, which did not overtly alter fibrillar collagen or the ECM proteome evaluated by quantitative mass spectrometry and N-terminomics. However, the distribution and quantity of collagen VI, a microfibrillar collagen that forms an open network with the basement membrane, was altered. In fibroblast-ablated mice, cardiac function was better preserved following angiotensin II/phenylephrine (AngII/PE)-induced fibrosis and myocardial infarction. Analysis of cardiomyocyte function demonstrated weaker contractions and slowed calcium decline in both uninjured and AngII/PE infused fibroblast-ablated mice. Moreover, fibroblast-ablated hearts had a similar gene expression profile to hearts with physiological hypertrophy after AngII/PE infusion. Our results indicate that the adult mouse heart tolerated a significant degree of fibroblast loss with potential beneficial impacts on cardiac function. Controlled fibroblast reduction may have therapeutic value in heart disease by providing cardioprotective effects.

## Introduction

Fibroblasts are considered to be the primary source of extracellular matrix (ECM) in the heart (1, 2). The main physiological role of the cardiac fibroblast is maintaining the ECM by balancing deposition and degradation of structural and nonstructural matricellular proteins (3). This ECM network serves as a scaffolding to mechanically support cardiomyocytes and permit transmission of lateral force by regulating mechanical signals (4, 5). It is thought that ECM-cytoskeletal connections are essential for proper stability and contraction of the cardiomyocyte - and disruptions in these interactions underlie a wide range of cardiomyopathies (6). Although the ECM network is considered to be primarily synthesized and organized by cardiac fibroblasts and integral to proper cardiomyocyte function (1, 2, 7), there is limited data focusing on homeostatic, fibroblast-specific ECM production *in vivo* and the potential impact of fibroblast reduction after the formation of the ECM scaffold but prior to injury.

In response to cardiac injury, fibroblasts rapidly adopt an activated phenotype resulting in their increased proliferation and deposition of a collagen-rich ECM (8-11). An initial adaptive response maintains structural integrity of the damaged myocardium and prevents rupture (12, 13). However, prolonged ECM deposition can impair cardiac compliance due to ventricular wall stiffening (14, 15). Persistent ECM accumulation resulting in fibrosis can also disrupt electrical transmission between cardiomyocytes leading to contractile dysfunction (16). Despite its clinical and pathophysiological significance, no interventions currently exist to directly treat or reverse cardiac fibrosis (17). Notwithstanding, the cardiac fibroblast has emerged as an ideal candidate to regulate fibrosis that accompanies cardiac injury, but its role remains relatively obscure.

Recent studies have focused on genetically targeting myofibroblasts or activated fibroblasts as a potential therapeutic treatment for heart disease because of their contribution to fibrotic scar formation and subsequent reduced heart function (10, 18-21). When chimeric antigen receptor (CAR) T cells were used to specifically target an endogenous cell-surface glycoprotein on activated fibroblasts, fibrosis was reduced leading to better cardiac function (22). Similarly, ablation of activated fibroblasts (21) or activated fibroblast-specific depletion of *Grk2*, a downstream effector of G protein-couple receptors that is known to be elevated in patients with heart failure (23), was shown to be cardioprotective after injury. While these studies suggest beneficial outcomes from fibroblast reduction, others have observed increased lethality and ventricular wall rupture, presumably due to reduced ECM deposition (18, 20), which was also observed when *Hsp47 (19), Fstl1 (24, 25)*, or *Smad3* (26) were disrupted in activated fibroblasts. Taken together, these data suggest that manipulation of fibroblast numbers and matrix deposition may require a more nuanced approach based on improved fundamental knowledge. Considering that anti-fibrotic therapies are currently being proposed as treatments for heart pathologies, a thorough evaluation and understanding of fibroblast activities during tissue homeostasis is necessary.

Given the lack of data specifically addressing fibroblast functions in the uninjured heart, we reduced fibroblast numbers by 60-80% in the adult murine heart using an inducible fibroblast-specific Cre line. Surprisingly, no mortality resulted even though fibroblast loss up to 80% was sustained 7 months post-induction. Moreover, heart function, protein composition, and proteolytic turnover were largely well-maintained despite the level of fibroblast depletion. Analysis of matrix components, such as type I, IV, and VI collagen demonstrated that the type I collagen fibrillar network appeared relatively unchanged, whereas changes in microfibrillar collagen and basement membrane components were observed. Fibroblast-ablated hearts also showed improved function after MI and angiotensin II/phenylephrine (AngII/PE) infusion, suggesting that an underlying reduction in fibroblast numbers or cardiac adaptation to fewer fibroblasts may be protective. Our findings reveal that fibroblast reduction before injury does not negatively affect the heart and may provide beneficial effects when cardiac remodeling or repair are provoked by injury.

## Results

### Genetic ablation of fibroblasts during cardiac homeostasis

After heart injury, fibroblasts are essential for the generation of replacement fibrous ECM and multiple studies have suggested that disruption of fibroblast expansion may lead to wall rupture (18, 20). Although it is thought that fibroblasts are indispensable for maintaining ECM structure during cardiac homeostasis, we recently demonstrated that a loss of up to 50% of resident fibroblasts was inconsequential for the architecture and function of the heart (27). These results led us to investigate whether the heart is able to tolerate a further reduction in fibroblast numbers. As PDGFRα is expressed in a majority of murine adult cardiac fibroblasts (8, 28), we induced diphtheria toxin A (DTA) expression using a *PDGFRα-CreER*^*T2/+*^ mouse line to deplete resident fibroblasts (29). After Cre induction by tamoxifen, DTA expression led to fibroblast loss within several days. We refer to these mice as fibroblast-ablated or ablated.

A tdTomato Cre recombination reporter demonstrated that a 60-80% reduction in PDGFRα-expressing cells occurred by 14 days post-induction (Figure 1A, D). To investigate fibroblast reduction independent of Cre expression, we assayed for vimentin expression (30) and found fewer vimentin-positive cells in ablated hearts compared to controls (Figure 1B). To determine whether other cell populations were capable of replacing fibroblast collagen production, we assessed cells actively expressing type I collagen using a transgenic collagen reporter, collagen1a1-GFP (*Col1a1*-GFP). This reporter expresses GFP under the control of the *Col1a1* promoter (10, 28, 31-33). A similar reduction in fibroblasts and collagen-expressing cells was observed (Figure 1C, E, F), suggesting that fibroblasts are the primary producers of type I collagen. Reduced reporter activity was not limited to the ventricles, as reductions were also observed in the atria and septum, while aortic expression of the GFP reporter remained relatively constant. To investigate the possibility that the remaining 20-40% of fibroblasts expanded and replenished fibroblast-ablated hearts over time, we quantified PDGFRα expression at 1 month and 7 months after induction. PDGFRα levels remained reduced in ablated hearts, suggesting that compensatory proliferation by residual fibroblasts does not occur (Figure 1I). Western blot, immunostaining, and qPCR confirmed a decrease in PDGFRα protein and transcript in ablated hearts 2-4 months post-induction (Figure 1G-J). Additionally, fibroblast related genes, including *Tcf21, Vim, Col1a1*, and *Col1a2*, were reduced in ablated hearts (Figure 1J). Furthermore, alternative cardiac fibroblast-specific genes identified in recent single cell RNAseq studies, including *Dpep1, Dkk3, Col5a1, Mdk*, and *Pdgfrl* (34, 35) were also reduced, suggesting that another population of fibroblasts does not compensate for the loss of PDGFRα expressing fibroblasts (Figure 1J). Flow cytometry confirmed the reduction in fibroblast numbers, while the number of endothelial (CD31^+^) and myeloid cells (CD11b^+^) were unaffected (Figure 1K). These data demonstrate that genetic fibroblast ablation is efficient and effectively reduces the resident fibroblast population within 14 days and lasts up to 7 months post-induction in the adult murine heart.

**Figure 1.**
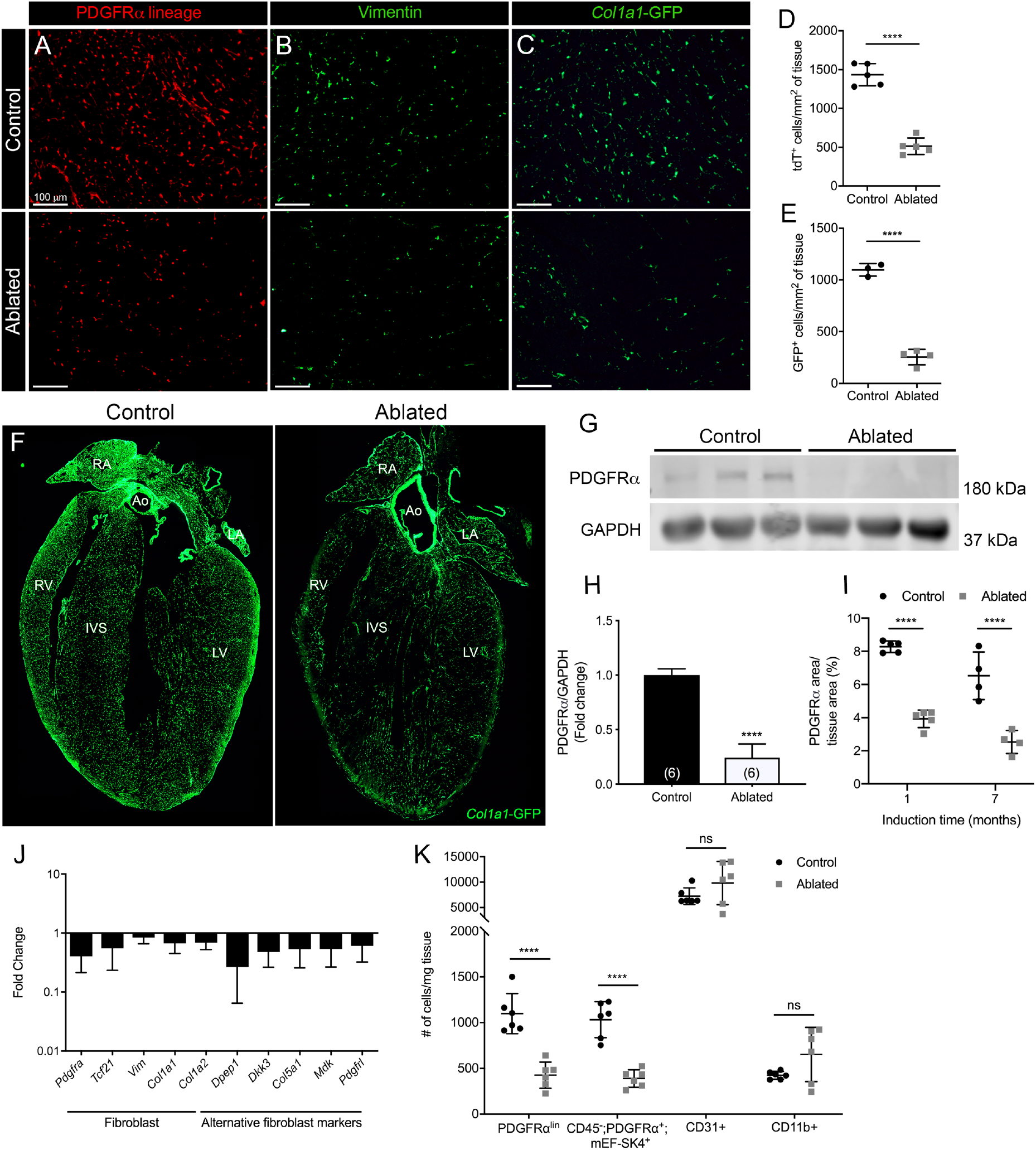
Loss of fibroblasts in adult ablated hearts. (**A-C**) Representative images of (**A**) PDGFRα lineage (tdTomato), (**B**) vimentin immunostaining, and (**C**) *Col1a1*-GFP reporter fluorescence in the left ventricular (LV) myocardium. Quantification of (**D**) tdTomato and (**E**) *Col1a1*-GFP. n=3-5 per group. (**F**) 4 chamber view of *Col1a1*-GFP expression. Ao: aorta, IVS: interventricular septum, LA: left atrium, LV: left ventricle, RA: right atrium, RV: right ventricle. (**G-H**) Representative western blots of whole ventricle lysates. n=6 per group. (**I**) Area of PDGFRα staining normalized to tissue area at indicated induction times. (**J**) qPCR analysis of selected fibroblast and alternative fibroblast genes in whole ventricle tissue from ablated hearts compared to controls. *18s* was used as a housekeeping gene. n=3-4 per group. (**K**) Quantification of fibroblasts (PDGFRα^lin^, CD45^-^;PDGFRα^+^;mEF-SK4^+^), endothelial cells (CD31^+^), and immune cell (CD11b^+^) populations by flow cytometry. Neutrophils, mast cells, and eosinophils were fewer than 100 cells/mg tissue in both control and ablated hearts. (A-F, K) 4-14 days after induction. (G, H, J) 2-4 months after induction. Results are mean ± SD. Statistical significance was determined by unpaired t-test. ns: not significant, *P* > 0.05; \*\*\*\**P* ≤ 0.0001.

### Collagen fiber organization and basement membrane protein distribution

Despite a significant reduction in fibroblasts, ablated hearts were histologically and functionally indistinguishable from controls. Although body weight was reduced in long-term fibroblast-ablated mice, heart weight to body weight ratio was proportionate in aged females and increased slightly in aged males (Figure 1–figure supplement 1A-C). Cardiomyocyte cross-sectional area (CSA) determined by wheat germ agglutinin (WGA) staining was similar in control and ablated hearts (Figure 1–figure supplement 1D). Echocardiography and pressure-volume loop analyses revealed that control and ablated hearts maintained a similar ejection fraction, left ventricular (LV) chamber size, and systolic and diastolic blood pressure change up to 7 months following fibroblast ablation (Figure 1–figure supplement 1E-H). Taken together these data signify that fibroblast loss is well tolerated in the heart in the absence of injury.

Because fibroblasts are responsible for the majority of type I collagen production in the heart (1, 2), we evaluated collagen levels by immunostaining and hydroxyproline content and found that type I collagen and hydroxyproline quantity remained constant in ablated hearts (Figure 2A, E, I). To examine the ultrastructure of the ECM, we visualized decellularized LV tissue by scanning electron microscopy (SEM). The endomysial weaves of collagen fibrils surrounding individual muscle fibers were less dense and thinner 2 months post-fibroblast ablation (Figure 2J). These results indicate that after fibroblast ablation, the adult murine heart retained physiological collagen abundance with modestly affected fibrillar collagen organization.

**Figure 2.**
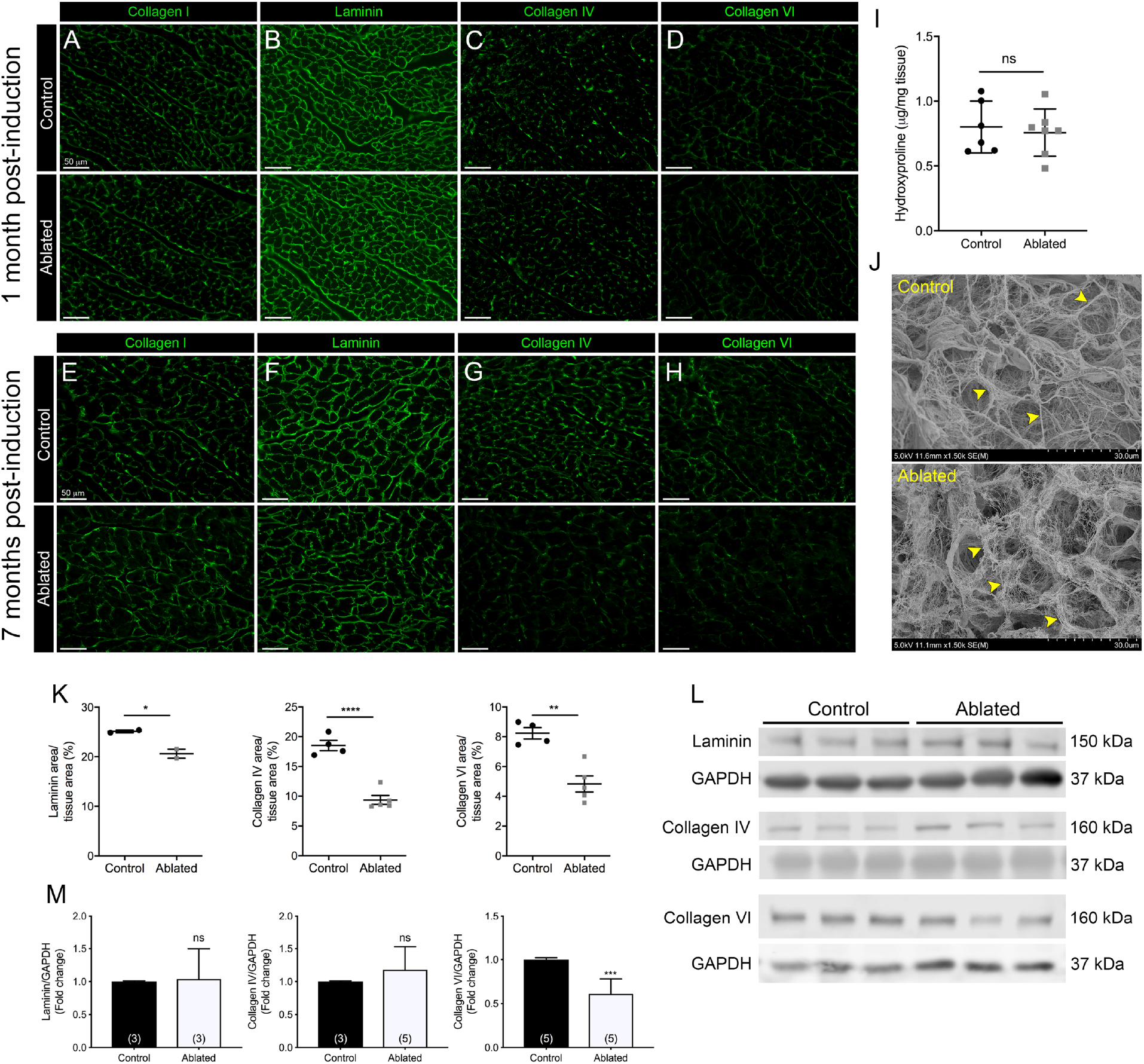
Collagen distribution and basement membrane alterations following fibroblast ablation. (**A-H**) Representative images of (**A, E**) collagen I, (**B, F**) laminin, (**C, G**) collagen IV, and (**D, H**) collagen VI immunostaining at indicated post-induction time points. (**I**) Hydroxyproline content from whole ventricle tissue 5 months after induction. (**J**) Representative scanning electron microscopy (SEM) images of decellularized LV tissue at 2 months post-induction. Arrowheads indicate collagen fibrils. (**K**) Area of laminin, collagen IV, and collagen VI staining normalized to tissue area 7 months after induction. (**L-M**) Western blot analysis of whole ventricle lysate 7 months after induction. n=3 per group. Results are mean ± SD. Statistical significance was determined by an unpaired t-test. ns: not significant, *P* > 0.05; \**P* ≤ 0.05; \*\**P* ≤ 0.01; \*\*\**P* ≤ 0.001; \*\*\*\**P* ≤ 0.0001.

Based on our previous finding showing that a 50% reduction in cardiac fibroblasts resulted in subtle alterations in laminin patterning, we hypothesized that the phenotype would be exacerbated with greater fibroblast loss (27). At 1 month post-induction, control and fibroblast-ablated hearts had similar patterns and levels of basement membrane proteins, including laminin, collagen IV, and collagen VI (Figure 2B-D). However, at 7 months post-induction, reductions in laminin, collagen IV, and collagen VI were observed by immunostaining in fibroblast-ablated hearts (Figure 2F-H, K). By western blot analysis, we observed a reduction in collagen VI but not laminin and collagen IV (Figure 2L-M), demonstrating that distribution of some basement membrane proteins, rather than extractable total protein, was affected by fibroblast loss (Figure 2L-M). These data suggest that loss of fibroblasts may disrupt basement membrane composition.

### Proteomics and degradomics of the ECM in fibroblast-ablated hearts

Fibroblasts are key in generating and remodeling many ECM components (36). Therefore, we evaluated cardiac protein composition using quantitative liquid chromatography-tandem mass spectrometry (LC-MS/MS)-based analysis of the decellularized ventricle to enrich for ECM proteins. The elicited cardiac proteomes showed only modest differences in composition (Figure 3A and Figure 3–figure supplement 1A), although principal component analysis and unsupervised hierarchical clustering demonstrated distinct segregation of decellularized proteomes of the control and fibroblast-ablated ventricles (Figure 3–figure supplement 1B-C). A statistical analysis of these proteomes identified several significantly altered secreted ECM proteins, including collagen VI α6 chain, collagen VIII α1 chain, chondroitin sulfate proteoglycan 4 (NG-2), and dermatopontin (Figure 3B and Figure 3–figure supplement 1C). The top affected pathways defined by Ingenuity Pathway Analysis (IPA) were GP6 signaling, apelin liver signaling, and the intrinsic and extrinsic prothrombin activation pathways (Figure 3–figure supplement 1D). Network analysis of all identified ECM protein ratios in IPA identified a molecular network associated with tissue development, cardiovascular system development and function, and cell morphology (Figure 3C).

**Figure 3.**
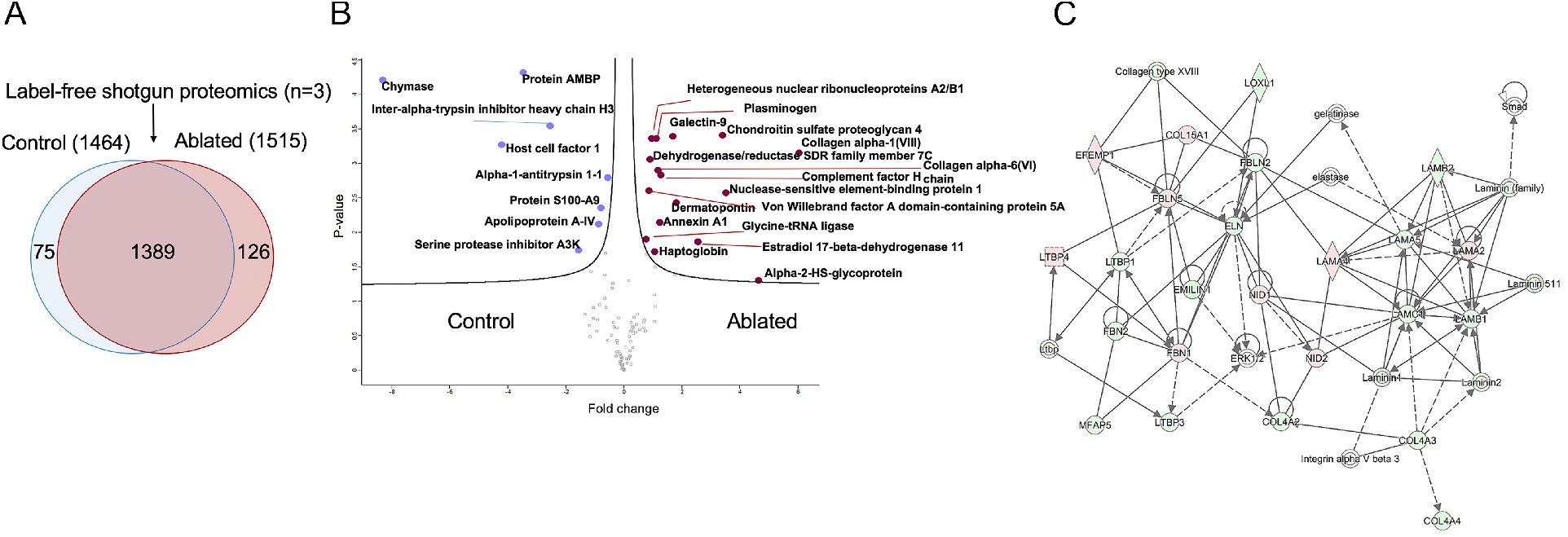
Minimal change in extracellular matrix (ECM) proteome after fibroblast ablation. (**A**) Venn diagram of proteins identified by label-free quantitative shotgun proteomics from decellularized control and ablated heart tissue >2 months after induction. (**B**) Volcano plot showing distribution of quantified proteins by shotgun proteomics according to p-value and fold change. Blue dots indicate proteins with higher abundance in control ventricles and red dots indicate proteins with higher abundance in fibroblast-ablated ventricles (all dots are FDR < 0.05). p-values are calculated from the data of three replicates. (**C**) ECM network generated using Ingenuity Pathway Analysis (IPA). Red and green nodes represent up-and down-regulated proteins in control samples, respectively. Empty nodes represent proteins that were not identified in this study but extrapolated from the IPA database. Dashed and continuous lines represent indirect and direct relationship between the proteins, respectively.

Using a specialized mass spectrometry-based approach, Terminal Amine Isotopic Labeling of Substrates (TAILS) (37), we identified changes in putative proteolytic events that occur in the reduction of fibroblasts using total protein from control and ablated hearts. TAILS is an N-terminomics method that specifically enriches for proteolytically cleaved (internal) peptides by labeling all N-termini at the protein level prior to trypsin digestion and LC-MS/MS (38). It thus enriches for peptides with labeled and/or naturally blocked N-termini (Figure 3–figure supplement 2A), whose origin and significance can be resolved bioinformatically (39), including determination of their location within the protein and the chemical moiety at the blocked/labeled N-terminus. These two determinations specify whether the observed N-terminus is a natural or internal one, and quantitative analysis determines which peptides are differentially abundant at a statistically significant level. We combined TAILS with LC-MS/MS analysis of peptides prior to enrichment and obtained 1834 blocked/labeled N-termini, including natural (signal peptide, signal peptide cleaved and modified, or termini arising from biosynthetic processing) and 1519 internal N-termini, which are likely to be generated by proteolysis (Figure 3–figure supplement 2B-C). Of these, 15 high confidence internal peptides corresponding to 8 proteins were more abundant in fibroblast-ablated hearts as compared to controls, implying higher proteolytic modification of these proteins, whereas 54 internal peptides corresponding to 30 proteins had lower abundance in fibroblast-ablated hearts (Supplemental Tables 1 and 2). The most internal peptides were identified in myosin-6 (Figure 3–figure supplement 3) and several were detected in other cellular proteins, especially actin isoforms, but no ECM-derived internal peptides were identified as statistically significant. Indeed, pathway analysis of the TAILS peptides identified an impact on cellular metabolism (e.g., mitochondrial dysfunction, oxidative phosphorylation, and sirtuin signaling) and the cytoskeleton and cell-cell adhesion (e.g., actin cytoskeleton and tight junction signaling), suggesting that cardiomyocyte remodeling may be occurring (Figure 3–figure supplement 2D).

### Resident fibroblast ablation mitigates cardiac impairment after myocardial infarction

Our data demonstrate that fibroblast loss does not lead to obvious structural or compositional deficiencies at baseline. However, depletion of activated fibroblasts after myocardial infarction (MI) has been shown to reduce collagen production and increase morbidity due to ventricular wall rupture (18). It is unclear how hearts with pre-existing resident fibroblast loss might respond to damage. To address this question, we induced MI by permanently ligating the left anterior descending (LAD) artery >7 weeks post-induction. At 10 weeks post-MI, ∼77% of control mice (7/9) and 100% of ablated mice (5/5) survived. Compared to controls, fibroblast ablation did not affect key parameters such as cardiac mass or lung weight, a measure of left-heart failure (Figure 4A-B). Echocardiography revealed that LV chamber size during diastole and systole were slightly reduced in ablated hearts after MI (Figure 4C-D). Examination of ejection fraction over time demonstrated that fibroblast-ablated mice had 5-20% improved ejection fraction compared to control mice (Figure 4E). However, cardiomyocyte CSA (Figure 4F) and the area of collagen (Figure 4G-H) remained similar between control and ablated hearts at 10 weeks post-MI. These results suggest that a 60-80% reduction in fibroblasts prior to injury does not eliminate replacement fibrosis during repair and ameliorates impairment of cardiac systolic function.

**Figure 4.**
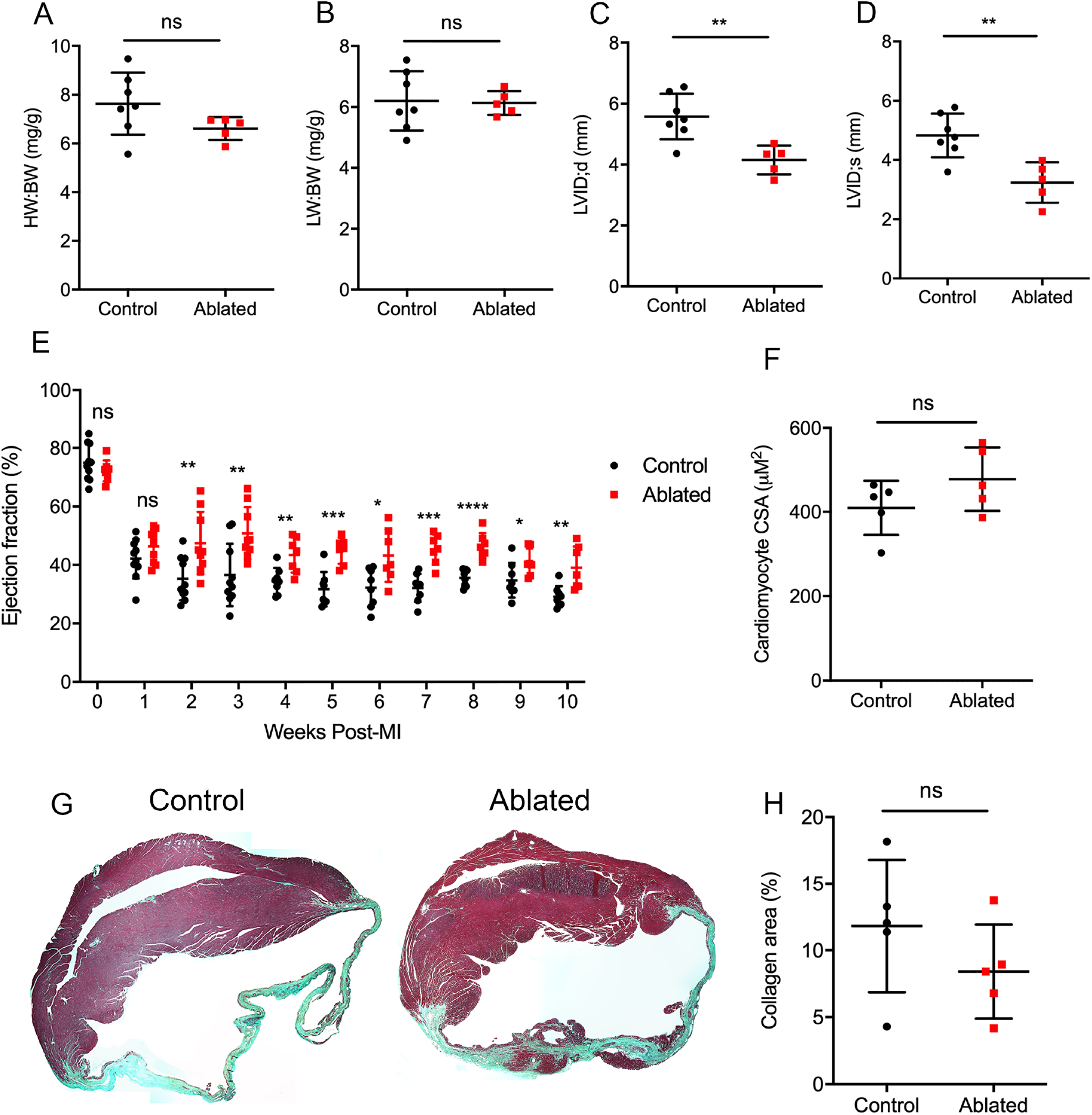
Key cardiac parameters and function after myocardial infarction (MI) in fibroblast-ablated mice. (**A**) Heart weight to body weight (HW:BW) ratio, (**B**) lung weight to body weight (LW:BW) ratio, (**C**) diastolic LV internal diameter (LVID), and (**D**) systolic LVID at 10 weeks post-MI. (**E**) LV ejection fraction (EF) post-MI. (**F**) Cardiomyocyte cross-sectional area (CSA) 10 weeks post-MI. (**G**) Representative images of trichrome staining and (**H**) quantification of percent collagen in control and ablated hearts. (A-G) >7 weeks post-induction at the time of ligation. Results are mean ± SD. EF results are mean ± SEM. Statistical significance was determined by an unpaired t-test. ns: not significant, *P* > 0.05; \**P* ≤ 0.05; \*\**P* ≤ 0.01; \*\*\**P* ≤ 0.001; \*\*\*\**P* ≤ 0.0001.

### Cardiac responses after Angiotensin II/phenylephrine infusion

The fibrotic response to injury is variable and disease-dependent (40). MI typically results in cardiomyocyte death and replacement fibrosis, while reactive fibrosis is thought to be induced by cardiac stress and inflammation (15). We, therefore, employed a second disease model to determine the impact of fibroblast loss on reactive fibrosis. We used angiotensin II and phenylephrine (AngII/PE) infusion to induce arterial hypertension, adaptive cardiac hypertrophy, and remodeling. Fibroblast levels remained relatively reduced as indicated by tdTomato expression (Figure 5–figure supplement 1). Fibroblasts activated by AngII/PE infusion demonstrated a perimysial pattern, surrounding cardiomyocytes. Similar to the response after MI, fibroblast ablation did not affect cardiac mass or lung weight (Figure 5A-B). Moreover, LV chamber size during diastole was not significantly changed but was slightly reduced during systole (Figure 5C-D). However, while control mice had a modest, sustained decrease in LV ejection fraction, the ejection fraction of hearts in ablated mice recovered to near baseline levels (Figure 5E).

**Figure 5.**
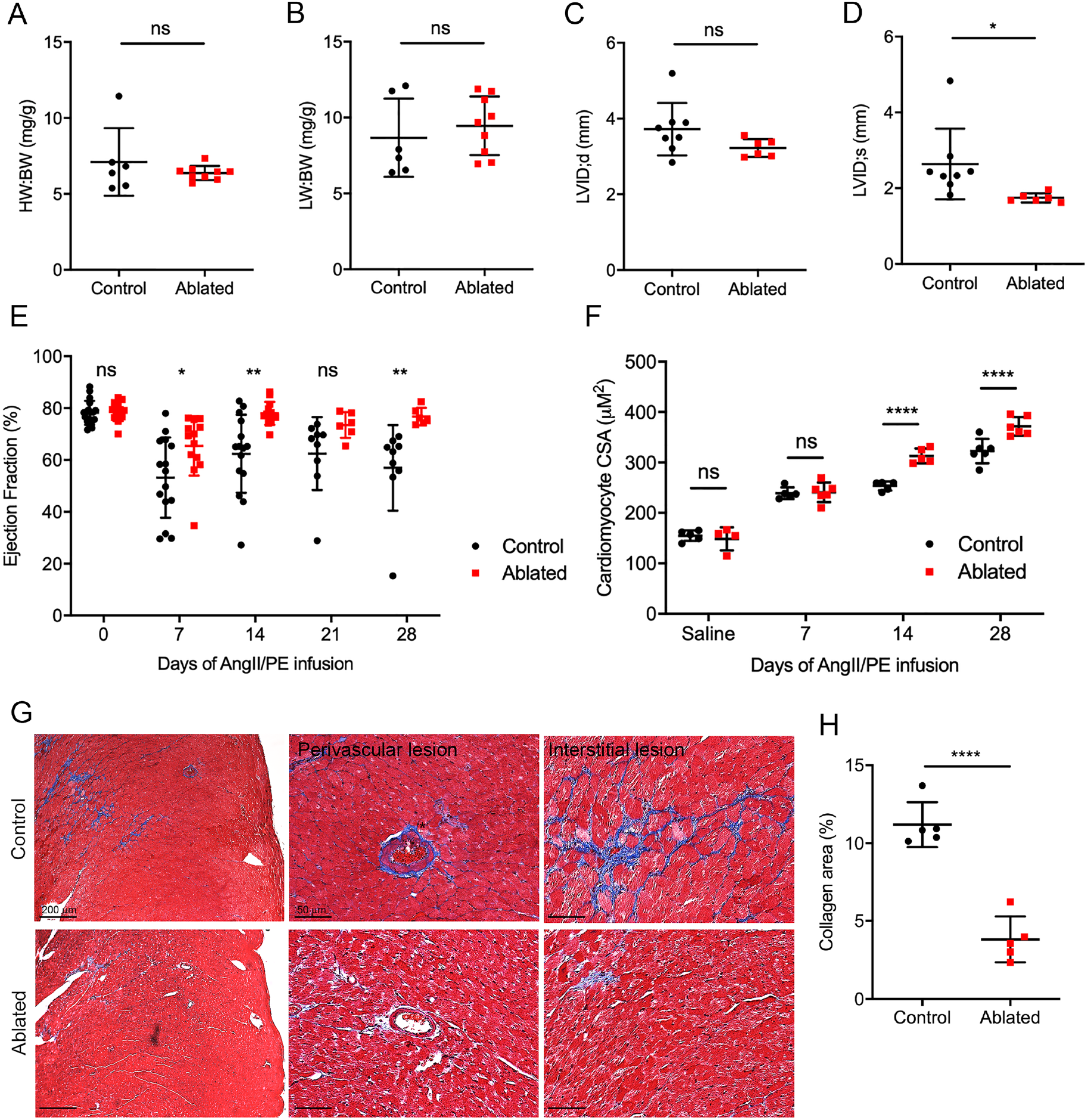
Effects of fibroblast loss on heart function and collagen accumulation after angiotensin II and phenylephrine (AngII/PE) infusion. (**A**) HW:BW ratio, (**B**) LW:BW ratio, (**C**) diastolic LVID, and (**D**) systolic LVID after 28 days of AngII/PE infusion. (**E**) EF and (**F**) cardiomyocyte CSA after AngII/PE infusion. (**G**) Representative images of trichrome staining of the LV myocardium showing perivascular and interstitial regions after 28 days of AngII/PE infusion. (**H**) Quantification of percent collagen in control and ablated hearts. (A-H) >7 weeks post-induction at the time of AngII/PE infusion. Results are mean ± SD. EF results are mean ± SEM. Statistical significance was determined by an unpaired t-test. ns: not significant, *P* > 0.05; \**P* ≤ 0.05; \*\**P* ≤ 0.01; \*\*\*\**P* ≤ 0.0001.

AngII stimulates hypertrophy in cardiomyocytes through angiotensin type 1 (AT1) receptors (41). While we did not observe an increase in heart to body weight ratio, we found an increase in cardiomyocyte CSA in AngII/PE treated, fibroblast-ablated hearts at both 14 and 28 days compared to control hearts (Figure 5F). We also observed a significant reduction in collagen deposition in adventitial and interstitial regions of the LV in fibroblast-ablated mice (Figure 5G-H). Taken together, these data suggest that ablating fibroblasts before injury results in greater cardiomyocyte hypertrophy and reduced fibrosis leading to normalization of ejection fraction.

### Gene expression profiling of fibroblast-ablated hearts after AngII/PE infusion

To identify potential molecular mechanisms that underlie the cardioprotective aspect of resident fibroblast loss during pathology, we identified differentially expressed genes by microarray analysis in fibroblast-ablated hearts compared to controls after 14 days of AngII/PE infusion. Forty-five genes were upregulated, and 204 genes were downregulated in ablated hearts (Figure 6–figure supplement 1A). Gene ontology (GO) analysis revealed that the downregulated genes were involved in cell adhesion, collagen binding, collagen fibril organization, and ECM organization (Figure 6A). The upregulated genes were found to be involved in the cellular response to extracellular stimulus and were localized to the membrane, mitochondria, and Z-disc (Figure 6A). Consistent with the observed reduction in fibrosis, we found decreased levels of numerous ECM-related mRNAs in ablated hearts by microarray analysis, including *Col1a1, Col1a2, Col3a1, Col6a2, Col6a3, Fn1, Tnc, Nid*, and *Fbn1* (Figure 6–figure supplement 1C). We also observed decreased levels in activated fibroblast genes by microarray and qPCR, including *Postn* and *Acta2* (Figure 6C and Figure 6–figure supplement 1C), suggesting that ablation of the resident fibroblast population also reduces the activated fibroblast population. One characterization of heart failure due to pathological hypertrophy is reactivation of the fetal gene program (42). However, *Nppa, Nppb*, and *Myh7* mRNAs were decreased in ablated hearts after 14 days of AngII/PE infusion, whereas *Myh6* was unchanged (Figure 6B-C), suggesting that pathological hypertrophy may not occur in ablated hearts. Because we observed cardiomyocyte hypertrophy in ablated hearts after AngII/PE infusion and a similar gene profile to physiological hypertrophy, we further examined genes differentially expressed in physiological hypertrophy, including *Insr* and *Igf1* (43), which revealed similar expression in ablated hearts (Figure 6B). qPCR validated key genes indicated by the microarray results (Figure 6C). Taken together, these data suggest that decreased fibroblast abundance not only led to reduced ECM gene expression but also elicited a potentially beneficial physiological cellular hypertrophic program.

**Figure 6.**
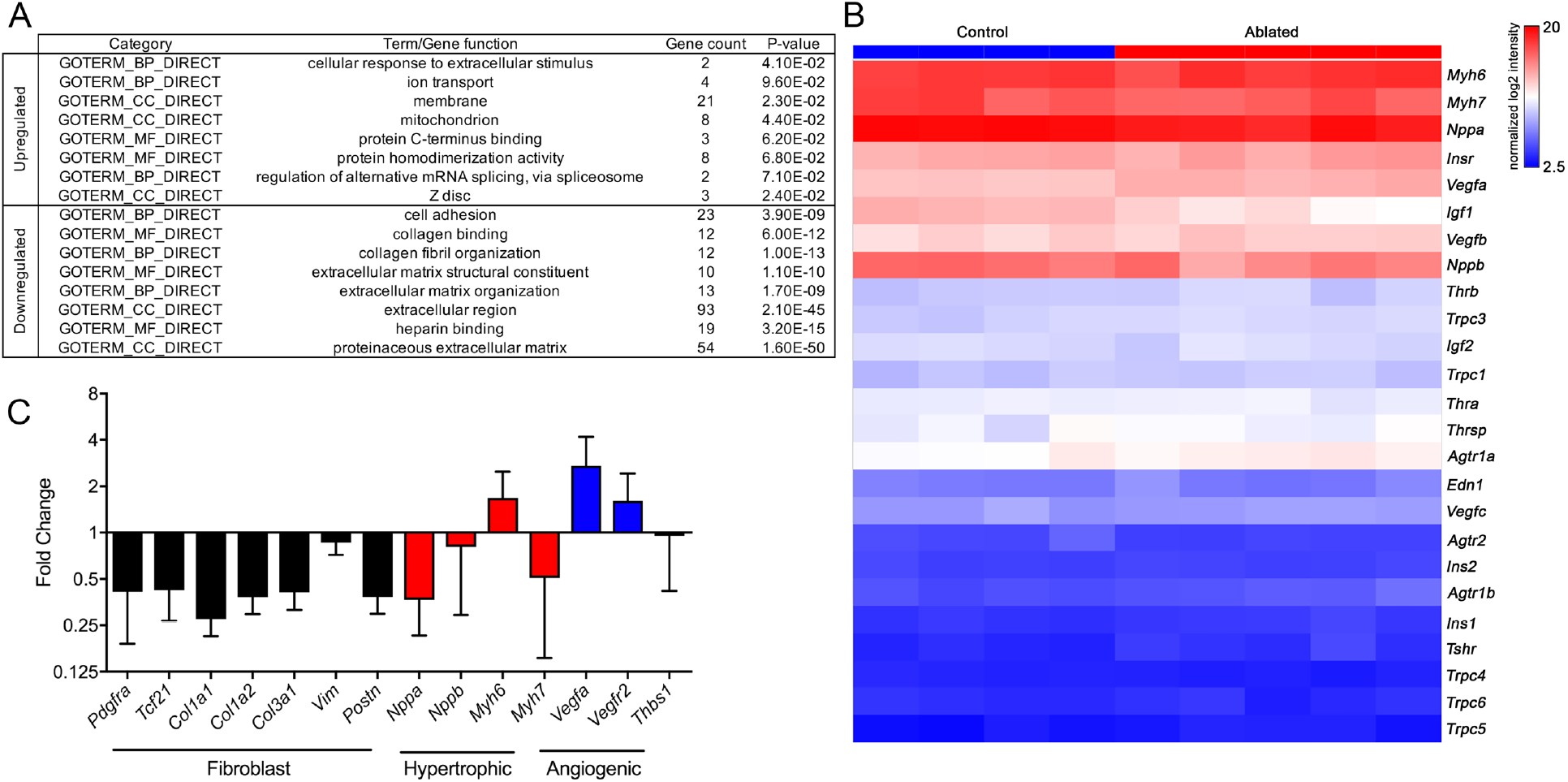
Gene expression profile after AngII/PE infusion. (**A**) Gene ontology (GO) analysis of differentially expressed genes between control and ablated hearts infused with AngII/PE for 14 days. All genes included had fold change values of log_2_ ≥ 2 or ≤ -2 and *P* ≤ 0.05. (**B**) Hierarchical clustering showing the expression levels of genes differentially expressed in 14 day AngII/PE infused control and ablated. n=4-5 per group. (**C**) qPCR analysis in whole ventricle tissue infused with 14 days of AngII/PE from ablated hearts compared to controls. GAPDH was used as a housekeeping gene. n=4-5 per group. Results are mean ± SD.

### Disruption of mechanical force generation and calcium reuptake in cardiomyocytes

Given the role of cardiomyocyte integrin signaling through interaction with basement membrane components (44, 45) and the observed changes in basement membrane composition of fibroblast-ablated hearts, we determined whether fibroblast loss led to alterations in cardiomyocyte function. Cardiomyocyte function was measured by recording contraction and calcium transients in individual cardiomyocytes. At baseline, we found that cardiomyocytes from ablated hearts had reduced sarcomeric shortening (Figure 7A), reduced speed of contraction (Figure 7B), and slower relaxation (Figure 7C), indicating decreased mechanical force. Differences were also observed in sarcomeric shortening and systolic velocity after 14 days of AngII/PE infusion (Figure 7A-B). Because calcium is a major determinant of cardiac contractility (46), we examined intracellular calcium handling in cardiomyocytes from fibroblast-ablated hearts. Calcium amplitude during contraction was similar between cardiomyocytes from control and ablated hearts at baseline and after AngII/PE infusion (Figure 7D). After AngII/PE infusion, calcium amplitude in cardiomyocytes increased from baseline, indicating the expected response to drug infusion (Figure 7D). Calcium reuptake, indicated by the decay time constant and the time to reach baseline, was slower at baseline and after AngII/PE infusion in fibroblast-ablated hearts (Figure 7E-F). These data suggest that the force of contraction and calcium efflux are deficient in cardiomyocytes when fibroblasts are depleted. Data from TAILS analysis support the observed cardiomyocyte deficiencies in ablated hearts by demonstrating that internal peptides from proteins involved in cardiomyocyte contraction, energy metabolism, and mitochondrial respiration were significantly changed in abundance in fibroblast-ablated hearts compared to controls. These included peptides from myosin-6, other myosins, several actin isoforms, and mitochondrial enzymes (Supplemental Tables 1 and 2). While these phenomena do not appear to negatively impact heart function, the weakened contraction and slowed calcium decline in cardiomyocytes may precondition and protect the heart during drug agonist induced-fibrosis.

**Figure 7.**
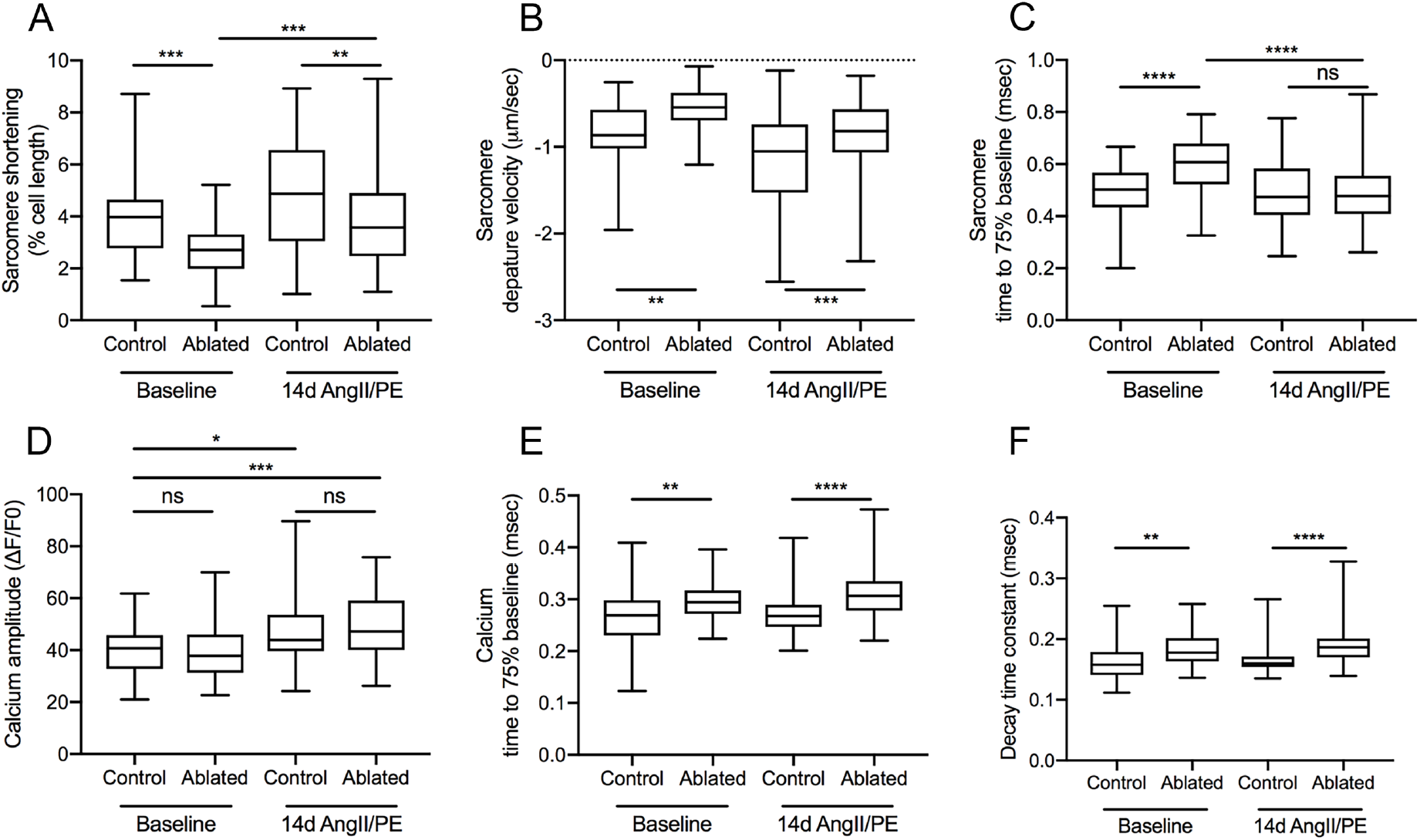
Cardiomyocyte contraction and calcium handling after fibroblast ablation. Cardiomyocyte (**A-C**) contraction and (**D-F**) calcium handling recording by edge detection from control and ablated hearts at baseline and after 14 days of AngII/PE infusion. >50 cardiomyocytes per group. n=5-6 per group, (A-F) 6-9 months after induction. Results are mean ± SD. Statistical significance was determined by an unpaired t-test. ns: not significant, *P* > 0.05; \**P* ≤ 0.05; \*\**P* ≤ 0.01; \*\*\**P* ≤ 0.001; \*\*\*\**P* ≤ 0.0001.

## Discussion

Our knowledge of fibroblast biology in the injured heart is rapidly expanding. Because fibroblast-targeting therapies are being contemplated as a mitigation strategy in fibrosis-associated cardiovascular diseases, an understanding of the homeostatic roles of fibroblasts is important. Fibroblasts are thought to be the primary source of type I collagen, and homeostatic collagen turnover and synthesis in the rat heart is estimated to be between 6-9% per day (47). Yet, our study reveals that the murine adult heart is functionally capable of sustaining a 60-80% reduction in fibroblasts for up to 7 months after ablation. Even though a complete ablation of fibroblasts was not obtained, the remaining fibroblasts did not expand and repopulate the heart. Despite this substantial loss of fibroblasts, proteomics of the decellularized ventricles did not detect massive changes in protein abundance, with type I collagen protein remaining relatively normal and fibrillar organization retained. Proteomics analysis is challenged by the abundance of cardiomyocyte components, post-translational modifications, and limited extraction of cross-linked components. Nonetheless, the collective data suggests that ECM in the unchallenged adult heart, once formed, is stable and has a low turnover rate, consistent with results from multi-isotope imaging mass spectrometry (MIMS), suggesting that ECM proteins have half-lives on the order of years (48).

The significant changes identified by TAILS in fibroblast-ablated hearts were surprisingly few in number and did not include any ECM-derived peptides, despite extensive offline fractionation prior to LC-MS/MS and identification of a large number of internal protein N-termini, suggesting that fibroblast depletion may not dramatically impact ECM proteolysis. One explanation is that although fibroblasts are primarily responsible for ECM degradation and deposition, there may be relatively low turnover at baseline in healthy hearts, and a 60-80% loss of these cells may not result in a significant change in protein composition. Alternatively, it is possible that the high sample complexity introduced by abundant cardiomyocyte proteins may have reduced the yield of ECM peptides, consistent with the finding that of the 1834 peptides, only 4.3% originated in secreted/extracellular matrix proteins. Nevertheless, this approach, in addition to LFQ-proteomics and cardiac function analysis, also supports a relatively mild impact of fibroblast ablation overall on the heart, although significant changes in several contractile proteins and metabolic enzymes were identified.

In support of previous findings (27), there was not a mechanism within the heart that signals fibroblast replenishment even with reductions up to 75%. It has been suggested that cardiac fibroblasts are heterogeneous (34, 35), and there is a possibility that a PDGFRα-negative fibroblast population compensates for fibroblast loss. However, analysis of total heart RNA argues that type I collagen transcript and reporter transgene levels remain reduced. Thus, in an undamaged heart, a loss in collagen synthesis might be balanced by an equal loss in collagen degradation, another action that has been primarily attributed to fibroblasts.

While severe changes to fibrillar collagen were not observed, there were more notable differences in the basement membrane and microfibrillar collagen surrounding cardiomyocytes. Therefore, these data suggest that fibroblasts produce factors, including matricellular proteins that may stabilize the fibrillar collagen network via cross-linking or proteolytic enzymes that regulate turnover of fibrillar collagens. Because type VI collagen is secreted by fibroblasts and connects the basement membrane to fibrillar collagen (49), we suspected that the observed basement membrane alterations in fibroblast-ablated hearts may be directly due to fibroblast loss. Alternatively, it is also possible that the observed basement membrane changes may be indirectly caused by disruption of collagen matrix organization or that the basement membrane and cell-proximate networks are not as well protected from turnover as cross-linked collagen. Collagen VI was reported to be overexpressed in hypertension, diabetes, and post-MI (49, 50). Interestingly, a recent study demonstrated that *Col6a1* knockout mice had improved heart function after MI (51). Therefore, our results are consistent with previous data in that the observed reduction in collagen VI at baseline and after 14 days of AngII/PE could, in part, explain the cardioprotective effect in fibroblast-ablated hearts after injury.

Because force generation is dependent on cardiomyocyte adhesion to the ECM which is mediated by the basement membrane, we expected to observe a disruption of cardiomyocyte contraction in fibroblast-ablated hearts. Indeed, our study is among the first to demonstrate that *in vivo* modulation of the ECM by fibroblasts affects cardiomyocyte function by abating contractile force and calcium efflux. One explanation is that the disrupted basement membrane in ablated hearts could decrease myofilament calcium sensitivity, which could be a result of reduced free calcium or force of contraction by the myofilaments (52). While we did not examine free calcium in our hearts, the disrupted basement membrane could alter myofilament adhesion leading to reduced contraction of the sarcomeres. Changes in contraction efficiency could also be a result of the observed altered proteolysis of proteins that are part of the contractile machinery, such as myosin-6. Thin filament proteins can undergo modifications, such as proteolysis leading to thin filament deactivation and slowed myocardial relaxation (53). In response to AngII/PE infusion, weaker contractions and slowed calcium decline in cardiomyocytes, as well as sustained normal cardiac function were also observed in fibroblast-ablated hearts. Therefore, we hypothesized that fibroblast loss predisposes the heart to physiological, rather than pathological hypertrophy in response to drug-induced fibrosis. However, because we used a relatively early time point for analysis of AngII/PE infusion, long-term injury models will be required to determine whether physiological hypertrophy is sustained over time in fibroblast-ablated hearts.

Because activated fibroblasts are key contributors to scar formation, fibroblasts have increasingly become a target of interest in combating heart disease. However, studies that have focused on specifically reducing activated fibroblasts or targeting genes in activated fibroblasts to attenuate pathological fibrosis have produced conflicting results. In some circumstances, fibroblast disruption leads to rupture while in other scenarios manipulations of fibroblasts appears to be protective (18-26). Our work further demonstrates that resident fibroblast reduction prior to injury may have beneficial outcomes under certain pathological conditions. Similarly, depletion of the resident fibroblast population prior to AngII/PE infusion reduced reactive fibrosis, however replacement fibrosis was not affected after MI. The minimal scar in the MI model could explain why depletion of fibroblasts prior to injury does not result in rupture of the ventricular wall, in contrast to what others have reported (18, 20). Because we only observe a 60-80% deletion of fibroblasts, the remaining 20-40% may be able to maintain the damaged myocardium after MI. This suggests that lowering level of initial fibroblasts could have beneficial effects in pathological conditions particularly in response to reactive fibrosis.

While we did not evaluate the ECM protein composition of the fibrotic scar, we predict that the ECM prior to injury may be altered resulting in subtle changes in matrix organization and stiffness. Although our proteomics data did not detect large differences in ECM components of fibroblast-ablated hearts at baseline, the differentially abundant ECM proteins that were observed could contribute to the protective response after injury. Because deletion of fibroblasts occurs several weeks before injury, the heart may have adapted prior to insult. Moreover, fibroblast loss increased cardiomyocyte hypertrophy in response to injury implicating a dynamic interaction between fibroblast and cardiomyocytes during pathology. Further investigation of the cardioprotective effect of resident fibroblast depletion will provide insights into the potential efficacy of anti-fibrotic therapies and delineate the long-term effects on the ECM and cardiac function.

In summary, these studies demonstrate the surprising finding that a significant reduction in cardiac fibroblasts is not detrimental to basal heart function. These observations suggest that cardiac tissue and especially the ECM may be very resilient. Further studies to determine the level of fibroblast loss that can be tolerated by cardiac tissue are warranted. Fibroblast loss prior to injury potentially resulted in a type of pre-conditioning that may be cardioprotective after injury. Further examination of the effect of fibroblast ablation after injury will provide insight into whether manipulation of the fibroblasts at specific stages of the pathological response may also have therapeutic value. Our study reinforces the idea that controlled fibroblast reduction may be a potential strategy in reducing maladaptive fibrosis in heart failure and other sustained cardiac diseases.

## Methods

### Mice

All animal protocols and experiments were approved by the University of Hawaii at Manoa Institutional Animal Care and Use Committee. Both males and females on a mixed C57Bl/6 background were used for these studies. *R26R*^*tdT*^ (54) (Jackson labs, 007914) and *R26R*^*DTA*^ (55) (Jackson labs, 010527) mice were purchased from Jackson Laboratory. *Collagen1a1-GFP* transgenic mice express cytoplasmic GFP under the control of a *col1a1* promoter/enhancer and were generated by Dr. David Brenner (33). *PDGFRα*^*CreERT2/+*^ mice were kindly provided by Dr. Brigid Hogan (Duke University Medical Center) (29). DTA expression was induced between 8 to 10 weeks of age by oral gavage on two non-consecutive days using 0.2 mg/g of body weight of tamoxifen (AdipoGen, CDX-T0200). The genotype for fibroblast reduction was *PDGFRα*^*CreERT2/+*^; *R26R*^*tdT/DTA*^. All control mice used in these experiments were tamoxifen induced and age matched. Control genotypes were *PDGFRα*^*CreERT2/+*^; *R26R*^*tdT/+*^ or *PDGFRα*^*+/+*^; *R26R*^*tdT/DTA*^ (littermate controls). A mix of male and female littermate (Cre^-^) and non-littermate (Cre^+^) controls were used. All experiments were performed on adult mice older than 8 weeks.

### Screening for PDGFRα deletion

All mice used in these experiments were screened for fibroblast loss either by PDGFRα expression, *Col1a1GFP* transgene expression, or *PDGFRα* transcript levels in the heart or kidney. Kidney tissue was collected, stored in RNAlater Stabilization Solution (Invitrogen, AM7021), and processed for RNA extraction as described below. Mice with less than 45% reduction of either fibroblasts or *PDGFRα* expression were excluded from studies (Supplemental Table 3).

### Immunostaining and microscopy

Cardiac tissue was excised, washed with DPBS, and saturated with 3 M KCl to arrest the heart in diastole. Hearts were bisected coronally and fixed with freshly prepared 4% PFA for 2 h at room temperature or 10% neutral buffered formalin (NBF) for 24 h at 4°C. Tissue fixed with 4% PFA was cryoprotected and frozen embedded. Immunostaining was performed on 10 μm tissue cryosections as previously described (27). Primary antibodies used for immunostaining are listed in Supplemental Table 4. Primary antibodies were detected using secondary antibodies from Thermo Fisher at a 1:500 dilution for 1 h at room temperature. Nuclei were stained with DAPI (Roche, 10-236-276-001). Tissue fixed with 10% NBF was processed for paraffin embedding. Trichrome staining was performed on 5 μm tissue sections using Gomori’s Trichrome Stain Kit (VWR, 84000-308) according to the manufacturer’s protocol. A Zeiss Axiovert 200 microscope equipped with an Olympus DP71 camera was used for imaging. Images and figures were edited in Photoshop CS6 (Adobe).

### Western blot

Atria and valves were removed from isolated hearts. Whole ventricle tissue was homogenized in RIPA buffer with protease inhibitor cocktail (Bimake, B14001) using a Dounce homogenizer. Samples were centrifuged at 16,000 x g for 20 min at 4°C, and supernatant was collected. Blots were probed with primary antibody overnight at 4°C, and then incubated with the corresponding Li-Cor IRDye secondary antibody for 1 h at room temperature. Primary antibodies used for western blot are listed in Supplemental Table 4. An Odyssey CLx imaging system was used for detection and images were analyzed using Image Studio version 5.2.5 software (Li-Cor Biosciences).

### qRT-PCR

RNA isolation was performed on whole ventricle tissue using IBI Isolate DNA/RNA reagent (IBI Scientific, IB47602) and PureLink RNA mini kit (Thermo Fisher, 12183025) according to the manufacturer’s instructions. RNA quality and concentration were determined by spectrophotometry using a NanoDrop 2000 instrument (Thermo Fisher). cDNA was synthesized using M-MLV Reverse Transcriptase (Sigma, M1302) and random hexamer primer (Thermo Fisher, SO142). qPCR analysis was performed using PowerUp SYBR Green Master Mix (Thermo Fisher, A25742) and a LightCycler 480 instrument (Roche). Samples were run in triplicate and normalized to *18s* or *Gapdh* expression. The 2^-ΔΔCt^ method was used for determining relative gene expression levels. Primer sequences used for qRT-PCR are listed in Supplemental Table 5.

### Flow cytometry

Adult hearts were transcardially perfused with DPBS containing 0.9 mM Ca^2+^, and atria and valves removed. Single-cell suspensions were obtained as previously described (28). Briefly, tissue was minced and incubated with collagenase type IV (600 U/ml, Worthington, LS004188) and dispase II (1.2 U/ml, Thermo Fisher, 17105041) in DPBS containing 0.9 mM Ca^2+^ for 45 min at 37°C and filtered through both a 40 μM and then a 30 μM filter to ensure a single-cell suspension. Yellow amine-reactive live/dead cell dye (Thermo Fisher, L34959) was used for live/dead discrimination. Cells were then incubated in Fc block (1:100, Tonbo Bioscience, 70-0161-U500) followed by indicated antibodies and fixed with 1% PFA in flow cytometry buffer. Primary antibodies used for flow cytometry are listed in Supplemental Table 4. Accucheck counting beads (Thermo Fisher, PCB-100) were added according to the manufacturer’s protocol prior to acquisition. Data were acquired on an LSRFortessa (BD Bioscience) and analyzed using FlowJo software version 10 (BD Bioscience). Cell counts were calculated according to the Accucheck counting bead manufacturer’s recommendations and normalized to the weight of individual ventricle tissue.

### Hydroxyproline assay

Hydroxyproline assay (Cell Biolabs, STA-675) was performed on whole ventricle tissue isolated from adult hearts. Samples were homogenized in distilled H_2_O and hydrolyzed in 6 N HCl for 20 h at 95°C. Hydrolyzed contents were processed according to manufacturer’s instructions. Absorbance of the supernatant was read using Molecular Devices SpectraMax M3 microplate reader at 540 nm wavelength.

### Scanning electron microscopy on decellularized tissue

LV tissue was cut into a 3×3×3 mm cube and fixed in 10% NBF for 24 h at room temperature. Fixed tissue was washed with distilled H_2_O and decellularized in 10% NaOH until tissue was clear. Decellularized tissue was washed in distilled H_2_O twice for 30 min, and then washed overnight at room temperature. Tissue was fixed with 4% tannic acid for 4 h, post-fixed with 1% osmium tetroxide in 0.1M sodium cacodylate, dehydrated through an ethanol series, and dried in a Tousimis Samdri-795 critical point dryer. Samples were mounted on aluminum stubs with double-stick carbon tape and coated with gold/palladium in a Hummer 6.2 sputter coater. Samples were viewed and digital images were acquired with a Hitachi S-4800 Field Emission Scanning Electron Microscope at an accelerating voltage of 5kV.

### Decellularization of cardiac tissue for mass spectrometry

Whole ventricle tissue from adult mouse was cut into 8-10 uniform pieces, washed in distilled H_2_O with protease inhibitor for 30 min, and decellularized in 1% SDS in DPBS with protease inhibitor until tissue was clear. Timing of decellularization was determined by DNA quantification using DAPI staining to confirm lack of cells in tissue samples. Decellularized cardiac tissue was washed in distilled H_2_O with protease inhibitor 3 times for 5 min at room temperature with light agitation, and then washed overnight to completely remove detergent. Decellularized tissue was placed in 200 μL of Protein Extraction Reagent Type 4 with protease inhibitor and sonicated 10 times for 10 sec. At least 100 μg of protein was sent to the University of Mississippi Medical Center for mass spectrometry analysis. Mass spectrometry experiments were performed blinded.

### Proteomics of decellularized tissue

Lysates from decellularized heart tissue were obtained as described above, and reduced, alkylated, and trypsin-digested into peptides. The peptides were cleaned using a Sep-Pak Vac C18 cartridge (Waters Corporation) and analyzed label-free by liquid chromatography-tandem mass spectrometry using a Q Exactive mass spectrometer (Thermo Fisher). A 15 cm Å∼ 75 μm C18 column (5 μm particles with 100 Å pore size) was used and the nano-UPLC ran at 300 nL/min with a 150 min linear acetonitrile gradient (from 5 to 35% B over 150 min; A = 0.2% formic acid in water; B = 0.2% formic acid in 90% acetonitrile). Tandem mass spectrometry (MS/MS) was set up with an exclusion of 25 s, and the samples were run with high-energy collisional dissociation fragmentation at normalized collision energy of 30% and an isolation width of 2 m/z. The resolution setting was 70,000 for target values of the MS at 1e6 ions and in MS2 at resolution setting of 17,500 for 1e5 ions. Mass spectrometry analyses were performed at the University of Mississippi Medical Center.

RAW files were analyzed using Proteome Discoverer 2.2 (Thermo Fisher). Precursor mass tolerance was set at 10ppm and fragment mass tolerance was set at 0.6Da. Dynamic modification was set to oxidation (+15.995 Da (M)) and static modification was set to carbamidomethyl (+57.021 Da (C)). Samples were searched against the reviewed mouse database downloaded from Uniprot (on November 2018, with 16977 sequences). A strict false discovery rate (FDR) of 1% was applied. Label-free quantification was done based on precursor ion intensity and normalization was done using the total peptide amount (from all peptides identified). Proteins were included only if they were identified by at least two high-confidence peptides.

For further visualizations such as multi-scatter plot, principle component analysis (PCA), volcano plot, and heat map, normalized abundances from all the samples were imported in Perseus 1.6.5.0. Values were log 2(x) transformed and data was filtered by excluding proteins which were not identified in at least 50% of the samples. The missing values for proteins present >50%, but not all samples were imputed from the normal distribution feature. Multi-scatter plot and PCA analysis were performed using all proteins, whereas only ECM proteins were used for generating the volcano plot and unsupervised hierarchical clustering.

### Heart degradomics using Terminal Amine Isotopic Labeling of Substrates (TAILS)

Protein extraction was performed using T-Per tissue protein extraction reagent (Thermo Fisher) with protease inhibitor cocktail (Roche) and prepared for TAILS as previously described (37). Briefly, heart tissue was homogenized in 0.5 mL of T-Per on ice and centrifuged, supernatant was collected in a fresh tube (T-Per extract) and 4M GuHCl containing a protease-inhibitor cocktail was added to the pellet and incubated further at 4°C for 24 h on a rotary shaker (GuHCl extract). All subsequent processing was done separately for T-Per and GuHCl extracts. Protein estimation was done using the Bradford assay and 200 µg of protein from each heart was denatured, reduced, and alkylated as per the iTRAQ labeling protocol (SciX). iTRAQ labels were reconstituted in DMSO and samples were incubated with iTRAQ labels for 2 h in the dark. Excess iTRAQ reagent was quenched by incubation with 100 mM ammonium bicarbonate for 30 min in the dark. After labeling, the respective extracts were combined (i.e. T-Per extracts combined with GuHCl extracts) for subsequent handling and MS analysis. After methanol-chloroform precipitation, the pellet was reconstituted in 100 µl of 100 mM NaOH, 50 µl of H2O, and 100 µl of 100 mM HEPES. Samples were digested overnight with trypsin (1:50 ratio) at 37°C. Peptide quantitation was performed using Pierce Quantitative Colorimetric Peptide Assay Kit (Thermo Fisher, 23275) and 500 µg of each sample was retained for analysis by MS (pre-TAILS sample) (Figure 3–figure supplement 2A). The remaining sample was enriched for blocked N-Termini using ALD-HPG polymer to bind reactive (tryptic) N-termini (TAILS). Pre-TAILS and TAILS samples were fractionated by reverse phase high performance liquid chromatography (HPLC). 32 fractions were collected which were pooled in bins to generate 4 final fractions for analysis on the Orbitrap Fusion Lumos mass spectrometer at the Lerner Research Institute Proteomics and Metabolomics Core. A small portion of unfractionated sample was also retained and analyzed on the mass spectrometer. C18 clean-up of samples was done prior to LC-MS/MS. Bioinformatics analysis was done and N-termini identified with a false discovery rate < 1% were annotated essentially as recently described (37).

### Myocardial infarction

Adult mice >22.0 g were subjected to MI as previously described (8). Briefly, a thoracotomy was performed between the third and fourth rib to expose the LAD artery. The LAD proximal artery was permanently ligated using a 7.0 silk suture. Ligation was confirmed by visualization of LV blanching and ST elevation on the electrocardiogram.

### Osmotic pump implantation

Adult mice >20.0 g were infused with AngII/PE to induce cardiac hypertrophy and fibrosis. Mice were anesthetized with 1-2% isoflurane and mini-osmotic pumps (Alzet, 2001, 2002, or 2004) were implanted subcutaneously. A combination of 1.5 μg/g/day angiotensin II (Calbiochem, 05-23-0101) and 50 μg/g/day phenylephrine hydrochloride (Sigma, P6126) or saline, was infused for 7, 14, or 28 days.

### Echocardiography

Echocardiography was performed using a Vevo 2100 system (VisualSonics) to analyze cardiac function after MI and AngII/PE infusion in conscious mice. Briefly, hair removal cream was applied to the chest until all fur was removed from the area. A layer of ultrasound gel was applied to the animal’s chest, and the probe was lowered at the parasternal line, 90° between the probe and the heart. B- and M-modes were performed to record 2-dimensional and 1-dimensional transverse cardiac measurements and used to analyze LV function. Independent echocardiography on mice 7 months post-induction at The Ohio State University also found no differences in cardiac measurements between control and fibroblast-ablated hearts.

### Pressure-volume (PV) loop

Cardiac hemodynamic measurements were assessed via a closed chest approach using a 1.4 rodent pressure volume catheter (Transonic) advanced into the left ventricle through the right carotid artery (56). In brief, mice were anaesthetized by ketamine (55 mg kg^-1^) plus xylazine (15 mg kg^-1^) in saline solution and placed in supine position on a heat pad. Following a midline neck incision, the underlying muscles were pulled to expose the carotid artery. Using a 4-0 suture, the artery was tied and the pressure–volume catheter was advanced through the artery into the left ventricle of the heart. After 5–10 min of stabilization, values at baseline and stimulation at varying frequencies (4–10 Hz) were recorded. To measure the beta-adrenergic response, 5mg kg^-1^ dobutamine was injected intraperitoneal. All the measurement and analysis were performed on LabChart7 (AD Instruments).

### Cardiomyocyte cross-sectional area (CSA) quantification

Cardiomyocyte boundaries were identified by wheat germ agglutinin labelling in 10 μm tissue sections that were processed as described above. Cardiomyocytes in cross-section were defined by having a circular to oval shape, surrounded by circular capillaries. A total of 100 cardiomyocytes in a section were outlined and CSA was calculated using ImageJ (NIH).

### Cardiomyocyte isolation

Cardiomyocytes from whole ventricle tissue were isolated from adult mice using a modified Langendorff-free collagenase digestion protocol (57). Briefly, mice were anesthetized with an intraperitoneal (IP) injection of tribromoethanol (0.4 mg/g of body weight), the thoracic cavity was opened, and descending aorta was severed. The right ventricle was immediately flushed with EDTA buffer and the ascending aorta was clamped. The heart was excised and the left ventricle was perfused with EDTA and perfusion buffer. The heart was enzymatically digested with 0.5 mg/mL Collagenase Type II (Worthington, LS004176), 0.5 mg/mL Collagenase Type IV (Worthington, LS004188), and 0.05 mg/mL Protease Type XIV (Sigma, P5147). The atria and valves were removed and tissue was teased apart. Cells were dissociated by gentle trituration with a wide-bore pipette tip and cell suspension was filtered through a 100 μM nylon strainer. Cells were allowed to settle by gravity for 20 min, the supernatant was removed, and the cell pellet was resuspended with Tyrode’s solution (130 mM NaCl, 5 mM KCl, 10 mM HEPES, 10 mM glucose, 0.5 mM MgCl_2_, 1.2 mM CaCl_2_, pH 7.4) for Ca^2+^ transients and contraction measurements.

### Measurement of cardiomyocyte Ca^2+^ transients

Cardiomyocytes were loaded with 2 μM Fura-2 (Thermo Fisher, F1201) in Tyrode’s solution for 15 min. Cells were electrically paced at 1 Hz using and IonOptix Myopacer. Measurements were performed using IonWizard version 6.1 software (IonOptix). Images were obtained by a Nikon Eclipse TE2000-U camera. Amplitude of intracellular Ca^2+^ transient was calculated as the difference between peak and diastolic Ca^2+^ levels according to the equation (F-F0)/F0 after subtraction of background fluorescence. The kinetics of Ca^2+^ transient time constants were determined using exponential curve fitting. Measurements were made in more than 50 cardiomyocytes for each group from n=5-6 mice.

### Microarray analysis

Total RNA was isolated as described above. The quality of RNA was assessed by an Agilent 2100 Bioanalyzer and samples with an RNA integrity number value ≥8.0 were used for microarray analysis. mRNA transcription profile was determined by Clariom S assay for mouse (Thermo Fisher, 902931). Analysis was performed using Transcriptome Analysis Console software version 4.0.1. The differentially expressed genes with a log_2_ (fold change) ≥ 2 or ≤ -2 and *P* ≤ 0.05 were analyzed by the functional annotation tool in the Database for Annotation, Visualization and Integrated Discovery (DAVID) version 6.8 program to search for enriched gene ontology (GO) terms.

### Statistical analyses

All statistical analyses were conducted using Prism 8 (Graphpad Software, La Jolla, USA). Statistical analyses between two groups were analyzed using unpaired student’s t-test. Comparisons between groups were analyzed using one-way ANOVA with Tukey test. Statistical variability is expressed as mean ± SD or SEM as stated in figure legends; ns: not significant, *P* > 0.05; \**P* ≤ 0.05; \*\**P* ≤ 0.01; \*\*\**P* ≤ 0.001; \*\*\*\**P* ≤ 0.0001.

### Study approval

All mouse experiments were performed according to the animal experimental guidelines issued and approved by Institutional Animal Care and Use Committee of the University of Hawaii at Manoa and The Ohio State University Wexner Medical Center.

### Data availability

Jill T. Kuwabara, Sumit Bhutada, Vikram Shettigar, Greg S. Gojanovich, Lydia P. DeAngelo, Jack R. Heckl, Julia R. Jahansooz, Dillon K. Tacdol, Mark T. Ziolo, Suneel S. Apte, Michelle D. Tallquist. (2020) Consequences of fibroblast ablation in adult murine hearts. The mass spectrometry proteomics data have been deposited to the ProteomeXchange Consortium via the PRIDE partner repository with the dataset identifier PXD021741 (shotgun proteomics) and PXD021739 (N-terminomics).

## Author contributions

JTK, SA, and MDT conceived the study, designed experiments, and wrote the manuscript. JTK, SB, GSG, LPD, JRH, JRJ, and DT conducted experiments and gathered data. JTK and SB performed statistical analyses. MZ and VS performed PV loop analyses. SB and SSA performed and analyzed mass spectrometry results.

## Acknowledgements

This work was supported by NIH HL074257 (MDT), NHLBI Institutional Cardiology Training Grant T32 HL115505 (JTK), and American Heart Association Grants PRE29630019 (JTK) and GRNT33660474 (MDT) and by the Allen Distinguished Investigator Program, through support made by The Paul G. Allen Frontiers Group and the American Heart Association (S.S.A). The Fusion Lumos Instrument at the Lerner Research Institute was purchased via an NIH shared instrument grant, 1S10OD023436-We thank S. Sebastian and C. Applegarth for their excellent technical support. We also thank the JABSOM Histology and Imaging Core supported by RCMI-BRIDGES G12 MD007601; the Center for Cardiovascular Research Animal Physiology Core supported by NIH grant P30 GM103341; the UH Cancer Center Genomics and Bioinformatics Shared Resource supported by the NCI Cancer Center Support Grant (CCSG) P30 CA071789; and the Pacific Biosciences Research Center’s Biological Electron Microscope Facility at the University of Hawaii at Manoa.

