## Supplemental files for "Consequences of PDGFRα^+^ fibroblast reduction in adult murine hearts"

SUPPLEMENTAL MATERIAL

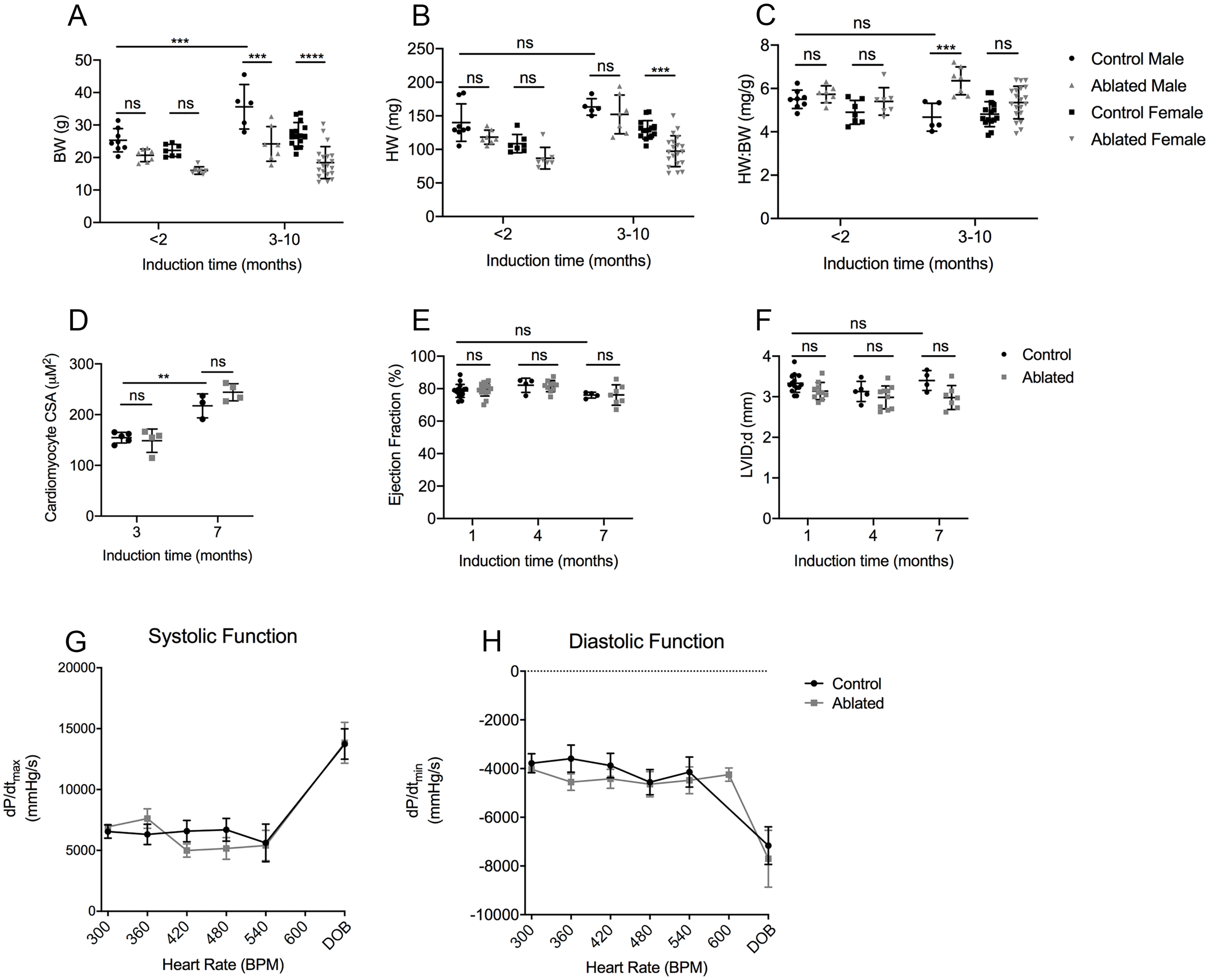


**Figure 1–figure supplement 1. Basal phenotype after short- and long-term fibroblast loss.** (**A**) BW, (**B**) HW, (**C**) HW:BW ratio, (**D**) cardiomyocyte CSA, (**E**) LV EF, and (**F**) diastolic LVID at indicated time of induction. (**G, H**) Pressure-volume loop analysis for systolic and diastolic function over increasing heart rate (HR). $dP/dt_{max}$ and $dP/dt_{min}$, maximum and minimum rate of pressure change in the ventricle, respectively. DOB: dobutamine. 7 months after induction. Results are mean ± SD. Statistical significance was determined by unpaired t-test. ns: not significant, *P* > 0.05; ***P* ≤ 0.01; ****P* ≤ 0.001; *****P* ≤ 0.0001.

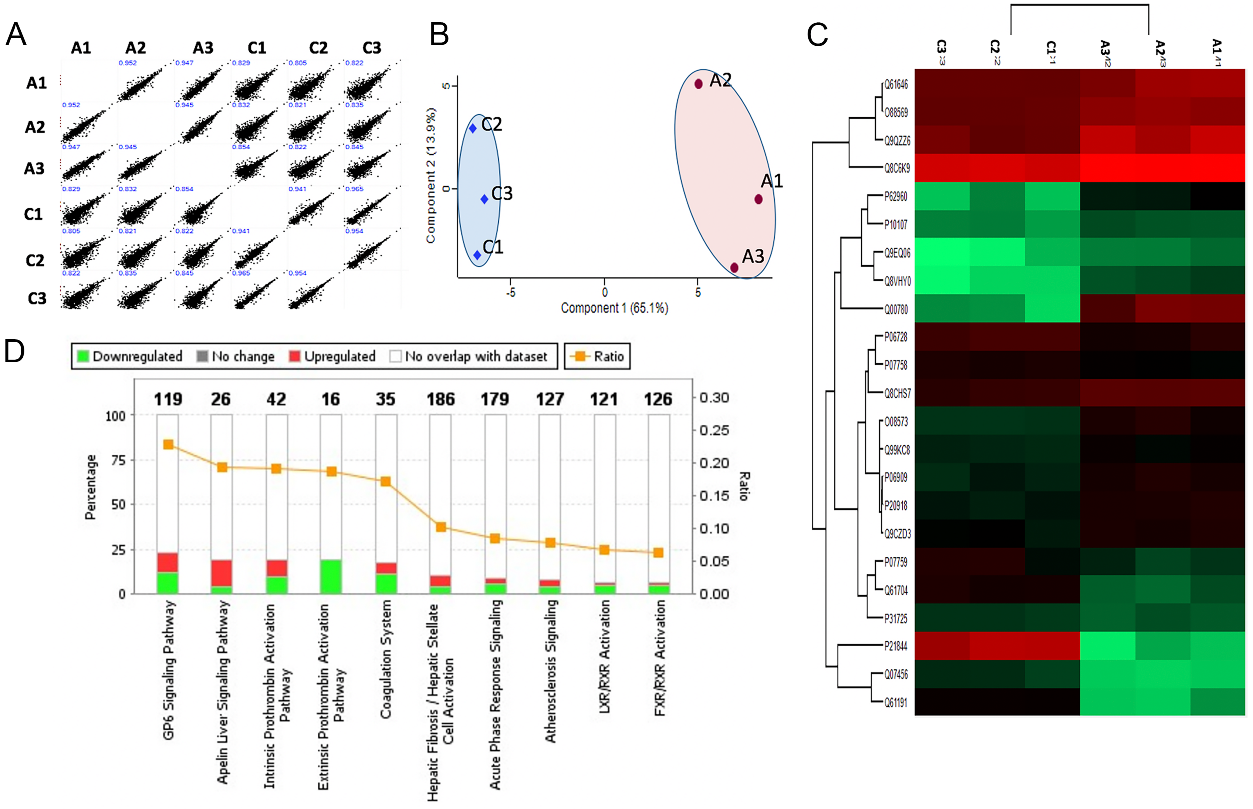


**Figure 3–figure supplement 1. Shotgun proteomics of decellularized heart tissue.** (**A**) Multi-sample scatter plots with Pearson's correlation coefficient value (blue text) showing strong correlation between individual samples. (**B**) Principal component analysis (PCA) of decellularized ventricular proteomes from control (C1, C2, C3) and ablated samples (A1, A2, A3). The first and second component segregation account for 65.1% and 13.9% of the differences between the two clusters, respectively. (**C**) Heat map of differentially abundant ECM proteins after unsupervised hierarchical clustering. (**D**) Top ten canonical pathways from secreted/ECM proteins showing statistically significant differential abundance.

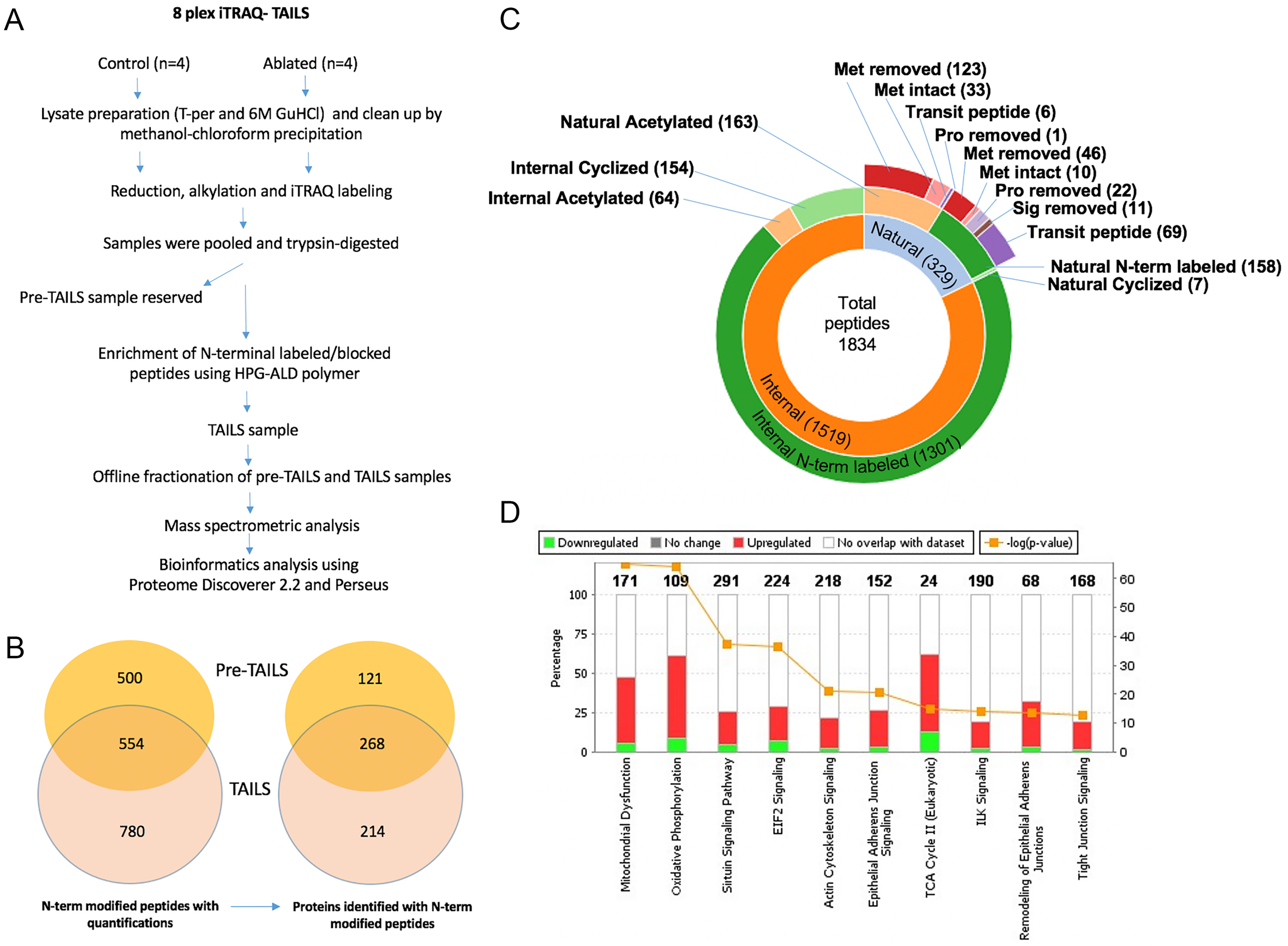


**Figure 3–figure supplement 2. N-Terminomics of whole ventricle tissue.** (**A**) Proteolytic changes were determined by 8-plex iTRAQ-TAILS N-terminomics on whole heart tissue using the illustrated workflow. (**B**) A Venn diagram illustrating the overall yield of N-terminally blocked/labeled peptides from the pre-TAILS and TAILS samples and the corresponding number of proteins they originated from. Of the total of 1834 N-terminally blocked/labeled peptides, 780 were identified exclusively in TAILS demonstrating the value of excluding unblocked tryptic peptides by HPG-ALD polymer. (**C**) Sunburst plot displaying all N-termini from fibroblast-ablated and control ventricles. The plot shows the distribution of blocked natural N-termini and internal N-termini in the innermost ring. The middle ring shows that the majority of internal N-termini were experimentally labeled with iTRAQ, whereas only a minority were acetylated or cyclized, contrasting with natural N-termini. The outermost ring indicates the origins of the natural N-termini. Met: methionine, Pro: propeptide, Sig: signal peptide. (**D**) Pathway analysis showing the top 10 canonical pathways differentially affected by fibroblast-ablation.

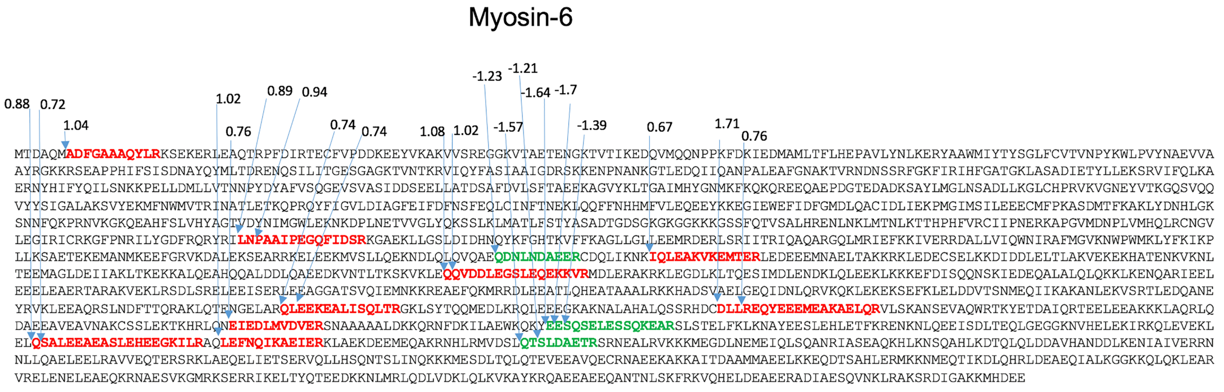


**Figure 3–figure supplement 3. Potential proteolytically cleaved peptides mapped on myosin-6.** Sequence of myosin-6 showing the locations of the internal peptides identified and indicating the respective increase (red peptides) or decrease (green peptides) in $log_2$ ratios of control/ablated levels. See also Supplemental Tables 1 and 2.

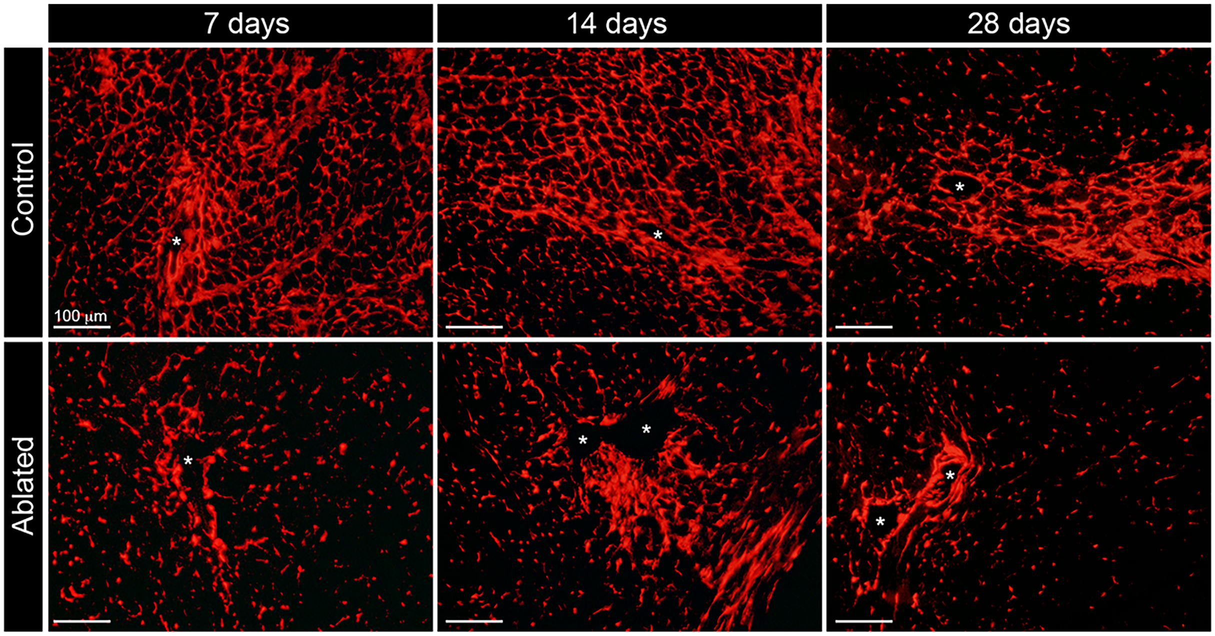


**Figure 5–figure supplement 1. Fibroblast expansion after AngII/PE infusion.** Representative images of tdTomato in the LV myocardium at 7, 14, and 28 days of AngII/PE infusion. Asterisk indicate blood vessel. n=3 per group.

A

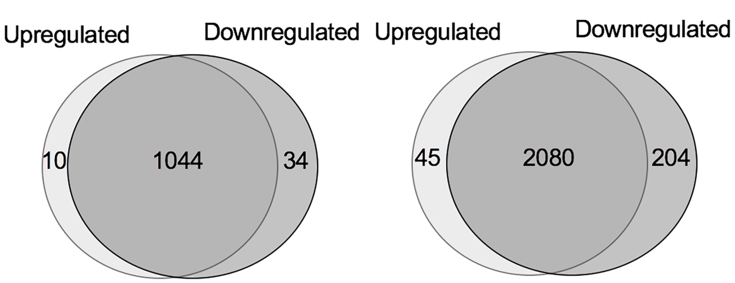


B

Baseline GO Analysis

| | Category | Term | Gene count | P-value |
|---|---|---|---|---|
| Upregulated | GOTERM_CC_DIRECT | external side of plasma membrane | 2 | 9.30E-02 |
| | GOTERM_CC_DIRECT | extracellular space | 3 | 7.10E-02 |
| Downregulated | GOTERM_BP_DIRECT | aging | 5 | 2.10E-04 |
| | GOTERM_MF_DIRECT | calcium ion binding | 6 | 4.60E-03 |
| | GOTERM_BP_DIRECT | cell adhesion | 5 | 9.10E-03 |
| | GOTERM_BP_DIRECT | extracellular fibril organization | 2 | 1.90E-02 |
| | GOTERM_CC_DIRECT | extracellular region | 22 | 4.00E-15 |
| | GOTERM_MF_DIRECT | heparin binding | 5 | 9.40E-05 |
| | GOTERM_BP_DIRECT | innate immune response | 6 | 5.50E-04 |
| | GOTERM_CC_DIRECT | proteinaceous extracellular matrix | 11 | 4.70E-11 |

C

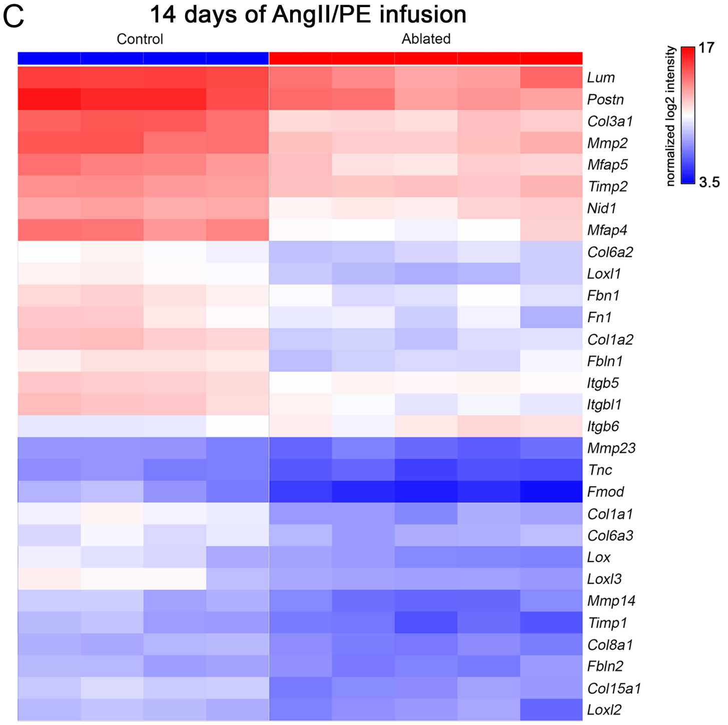


**Figure 6–figure supplement 1. Comparison of differentially expressed genes at baseline and after 14 days of AngII/PE infusion.** (**A**) Venn diagrams showing the number of upregulated and downregulated genes in ablated hearts compared to controls identified by microarray analysis. (**B**) GO analysis of differentially expressed genes between control and ablated hearts at baseline. (**C**) Hierarchical clustering showing the expression levels of ECM related genes differentially expressed between control and ablated hearts after 14 day AngII/PE infusion. n=4-5 per group. 7 weeks after induction. All genes included had fold change values of log2 ≥ 2 or ≤ -2 and $P \leq 0.05$.

**Supplemental Table 1. Potential proteolytically cleaved (internal) peptides with statistically significant higher abundance in control hearts.** The annotated sequence shows the cleaved peptide bond (.), the residue preceding the scissile bond (e.g., [A], corresponding to the P1 residue according to the nomenclature of Schechter and Berger (1). The position of the P1' residue (which follows the scissile bond) is presented in column 4.

| Accession number (UniProt) | Protein name | Annotated Sequence | P1' AA Position | (log2): (Control / Ablated) | Adj. P-Value: (Control / Ablated) |
|---|---|---|---|---|---|
| Q9CQ40 | 39S ribosomal protein L49, mitochondrial | [L].SQTQGPPDNPGFVESVDEYQFVER.[L] | 27 | 1.13 | 0.049233004 |
| Q99KI0 | Aconitate hydratase, mitochondrial | [R].YDLLEKNINIVR.[K] | 45 | 0.83 | 0.001630516 |
| P68134 | Actin, alpha skeletal muscle | [G].LVKAGFAGDDAPR.[A] | 18 | 1.65 | 6.23665E-06 |
| P60710 | Actin, cytoplasmic 1 | [W].DDMEKIWHHTFYNELR.[V] | 80 | 1.73 | 0.003692542 |
| P63268 | Actin, gamma-enteric smooth muscle | [NQ].EMATAASSSSLEKSYELPDGQVITIGNER.[F] | 226 | 1.04 | 4.9404E-05 |
| P63268 | Actin, gamma-enteric smooth muscle | [S].SSLEKSYELPDGQVITIGNER.[F]<br>[Y].VALDFENEMATAASSSSLEKSYELPDGQVITIGNER.[F]<br>[Y].ANNVLSGGTTMYPGIADR.[M]<br>[Q].GVMVGMGQKDSYVGDEAQSKR.[G]<br>[C].DIDIRKDLYANNVLSGGTTMYPGIADR.[M]<br>[S].SSSLEKSYELPDGQVITIGNER.[F]<br>[S].SLEKSYELPDGQVITIGNER.[F]<br>[S].LEKSYELPDGQVITIGNER.[F]<br>[G].FAGDDAPR.[A] | 234<br>221<br>297<br>42<br>288<br>233<br>235<br>236<br>21 | 0.88<br>0.83<br>1.14<br>0.85<br>2.22<br>0.87<br>0.71<br>0.75<br>0.94 | 0.00072102<br>0.001350125<br>0.001378523<br>0.001383431<br>0.003153518<br>0.003172322<br>0.007968922<br>0.012062644<br>0.012258745 |
| Q8K370 | Acyl-CoA dehydrogenase family member 10 | [H].STVAAASPSHEAKGGLVISPEGLSPAVR.[K] | 637 | 0.59 | 0.035970635 |
| P48962 | ADP/ATP translocase 1 | [C].FVYPLDFAR.[T] | 130 | 1.92 | 0.003905538 |
| P56480 | ATP synthase subunit beta, mitochondrial | [R].GVQKILQDYK.[S] | 423 | 0.58 | 0.042326665 |
| Q8BFZ3 | Beta-actin-like protein 2 | [Y].VAIQAVLSLYASGR.[T] | 134 | 1.53 | 0.045493203 |
| Q9CZU6 | Citrate synthase, mitochondrial | [H].ASASSTNLKDVLSNLIPKEQAR.[I] | 26 | 1.26 | 5.56119E-07 |
| Q68ED7 | CREB-regulated transcription coactivator 1 | [F].QSSGLDTSRTTR.[H] | 85 | 0.95 | 0.000204936 |
| Q9CZ13 | Cytochrome b-c1 complex subunit 1, mitochondrial | [T].ATFAQALQSVPETQVSILDNGLR.[V] | 36 | 0.84 | 0.003692542 |
| P19783 | Cytochrome c oxidase subunit 4 isoform 1, mitochondrial | [H].GSVVKSEDYAFPTYADRR.[D]<br>[A].HGSVVKSEDYAFPTYADRR.[D]<br>[G].SVVKSEDYAFPTYADRR.[D] | 25<br>24<br>26 | 1.45<br>1.31<br>1.27 | 1.28577E-09<br>1.11222E-06<br>0.000284487 |
| P12787 | Cytochrome c oxidase subunit 5A, mitochondrial | [H].GSHETDEEFDAR.[W] | 40 | 0.66 | 0.02951456 |
| Q9CPQ1 | Cytochrome c oxidase subunit 6C | [R].KAGIFQSAK.[-] | 68 | 1.49 | 1.28577E-09 |
| Q80XN0 | D-beta-hydroxybutyrate dehydrogenase, mitochondrial | [R].TTKSFLPLLR.[R] | 175 | 1.41 | 6.36704E-05 |
| Q9JKS4 | LIM domain-binding protein 3 | [Y].SAETLREMAQMYQMSLR.[G] | 200 | 0.7 | 0.030121229 |
| Q5SX40 | Myosin-1 | [V].AQWRTKYETDAIQR.[T]<br>[W].LPVYNAEVVAAYR.[G]<br>[L].PVYNAEVVAAYR.[G]<br>[P].VYNAEVVAAYR.[G] | 1372<br>131<br>132<br>133 | 1.82<br>1.2<br>1.13<br>0.99 | 0.003153518<br>0.008323708<br>0.010506035<br>0.041619738 |
| O08638 | Myosin-11 | [R].QKHTQAVEELTEQLEQFKR.[A] | 1199 | 0.59 | 0.042326665 |

| | | | | | |
|---|---|---|---|---|---|
| Q02566 | Myosin-6 | [E].QQVDDLEGSLEQEKKVR.[M]<br>[M].ADFGAAAQYLR.[K]<br>[Q].LEFNQIKAEIER.[K]<br>[I].LNPAAIPEGQFIDSR.[K]<br>[L].REQYEEEMEAKAELQR.[V]<br>[L].QSALEEAEASLEHEEGKILR.[A]<br>[Q].QVDDLEGSLEQEKKVR.[M]<br>[R].QLEEKEALISQLTR.[G]<br>[N].PAAIPEGQFIDSR.[K]<br>[Q].SALEEAEASLEHEEGKILR.[A]<br>[K].IQLEAKVKEMTER.[L]<br>[L].EEKEALISQLTR.[G]<br>[C].DLLREQYEEEMEAKAELQR.[V]<br>[Q].LEVEKLELQSALEEAEASLEHEEGKILR.[A]<br>[N].EIEDLMVDVER.[S] | 1029<br>7<br>1563<br>725<br>1346<br>1541<br>1030<br>1292<br>727<br>1542<br>913<br>1294<br>1343<br>1535<br>1424 | 1.08<br>1.04<br>0.9<br>0.89<br>0.76<br>0.88<br>1.02<br>0.74<br>0.94<br>0.72<br>0.67<br>0.74<br>1.71<br>1.02<br>0.76 | 5.39585E-05<br>0.000125178<br>0.001630516<br>0.004537537<br>0.006220997<br>0.006318734<br>0.009322742<br>0.010457201<br>0.010874433<br>0.013504196<br>0.015706071<br>0.023284843<br>0.027582409<br>0.033044627<br>0.039788277 |
| O70468 | Myosin-binding protein C, cardiac-type | [K].WLKDGVELTR.[E]<br>[C].STELFVKEPPVLITR.[S] | 482<br>440 | 0.68<br>1.23 | 0.01313844<br>0.016178058 |
| Q9D6J6 | NADH dehydrogenase [ubiquinone] flavoprotein 2, mitochondrial | [A].GGALFVHRDTPENNPDTPFDFTPENYKR.[I]<br>[A].GGALFVHR.[D] | 34<br>34 | 0.96<br>1.62 | 0.024431028<br>0.03218703 |
| P52503 | NADH dehydrogenase [ubiquinone] iron-sulfur protein 6, mitochondrial | [R].QKEVNENFAIDLIAQQPVNEVEHR.[I] | 52 | 0.88 | 0.001926476 |
| A2AAJ9 | Obscurin | [QR].MSKAAPVEWR.[K] | 4681 | 2.34 | 0.002949972 |
| O55126 | Protein NipSnap homolog 2 | [A].REDSWLKSLFVR.[K] | 36 | 1.78 | 1.73128E-12 |
| Q8K2B3 | Succinate dehydrogenase [ubiquinone] flavoprotein subunit, mitochondrial | [S].AKVSDAISTQYPVVDHEFDAVVVGAGGAGLR.[A] | 45 | 1.51 | 5.68118E-05 |
| P50752 | Troponin T, cardiac muscle | [F].MPNLVPPKIPDGERVDFDDIHR.[K] | 84 | 2.14 | 0.000261564 |
| P62984 | Ubiquitin-60S ribosomal protein L40 | [G].IIEPSLR.[Q] | 77 | 1.56 | 0.024771141 |
| P20152 | Vimentin | [Y].SSSPGGAYVTR.[S] | 54 | 1.22 | 7.1174E-07 |

**Supplemental Table 2. Potential proteolytically cleaved peptides with statistically significant higher abundance in fibroblast-ablated hearts.** The annotated sequence shows the cleaved peptide bond (.), the residue preceding the scissile bond (e.g., [A], corresponding to the P1 residue according to the nomenclature of Schecter and Berger (1). The position of the P1' residue (which follows the scissile bond) is presented in column 4.

| Accession number (UniProt) | Protein name | Annotated Sequence | P1' AA Position | (log2): (Control ) / (Ablated) | Adj. P-Value: (Control ) / (Ablated) |
|---|---|---|---|---|---|
| Q70FJ1 | A-kinase anchor protein 9 | [R].ELEQALLASAEPFP.[K] | 2115 | -2.03 | 0.006094 |
| Q5SX40 | Myosin-1 | [E].LEEEIEAER.[AT]<br>[Y].ETDAIQR.[T]<br>[L].LQAEIEELR.[A] | 1121<br>1379<br>1684 | -1.66<br>-1.15<br>-1.3 | 0.030995<br>0.001601<br>0.012259 |
| P13541 | Myosin-3 | [VI].EDLMVDVER.[AS] | 1426 | -1.35 | 0.037948 |
| Q5SX39 | Myosin-4 | [E].ATAAALR.[K] | 1190 | -1.29 | 0.000175 |
| Q02566 | Myosin-6 | [E].QDNLNDAEER.[C]<br>[Y].EESQSELESSQKEAR.[S]<br>[E].ESQSELESSQKEAR.[S]<br>[E].SQSELESSQKEAR.[S]<br>[L].QTSLDAETR.[S]<br>[ETA].SLDAETR.[RSA] | 897<br>1461<br>1462<br>1463<br>1598<br>1600 | -1.23<br>-1.64<br>-1.7<br>-1.39<br>-1.57<br>-1.21 | 0.014372<br>0.018789<br>0.000965<br>3.51E-05<br>2.69E-07<br>0.00071 |
| Q61941 | NAD(P) transhydrogenase, mitochondrial | [R].KTTVLAMDQVPR.[V] | 171 | -1.61 | 0.018789 |
| Q3V129 | Serine/threonine-protein kinase ULK4 | [D].PLPPIPKDSSFPK.[A] | 236 | -0.9 | 0.047896 |
| P58771 | Tropomyosin alpha-1 chain | [K].LKYKAISEELDHALNDMTSI. [-] | 265 | -2.64 | 4.71E-06 |

**Supplemental Table 3. Percent of PDGFRα deletion determined for each experiment.**

| Figure | Experiment | % deletion | Method |
|---|---|---|---|
| Figure 1A, 1D | tdTomato cell number quantification | 65 | tf |
| | | 72 | tf |
| | | 68 | tf |
| | | 64 | tf |
| | | 52 | tf |
| Figure 1C, 1E | GFP cell number quantification | 71 | gf |
| | | 87 | gf |
| | | 74 | gf |
| | | 75 | gf |
| Figure 1G-H | Western blot (2-4 months post-induction) | 79 | wb (PDGFRa) |
| | | 78 | wb (PDGFRa) |
| | | 93 | wb (PDGFRa) |
| | | 72 | wb (PDGFRa) |
| | | 79 | wb (PDGFRa) |
| | | 54 | wb (PDGFRa) |
| Figure 1I, Figure 2A-D | PDGFRa, collagen I, Laminin, collagen IV, collagen VI IHC (1 month post-induction) | 50 | ihc (PDGFRa) |
| | | 63 | ihc (PDGFRa) |
| | | 48 | ihc (PDGFRa) |
| | | 53 | ihc (PDGFRa) |
| | | 48 | ihc (PDGFRa) |
| | | all samples 50-70 | tf |
| Figure 1I, Figure 2E-H, 2K | PDGFRa, collagen I, Laminin, collagen IV, collagen VI IHC (7 month post-induction) | 60 | ihc (PDGFRa) |
| | | 57 | ihc (PDGFRa) |
| | | 47 | ihc (PDGFRa) |
| | | 74 | ihc (PDGFRa) |
| | | all samples 50-70 | tf |
| Figure 1J | Baseline qPCR | 61 | hq (PDGFRa) |
| | | 46 | hq (PDGFRa) |
| | | 72 | hq (PDGFRa) |
| Figure 1K | Flow cytometry | 70 | fc (PDGFRalin) |
| | | 80 | fc (PDGFRalin) |
| | | 42 | fc (PDGFRalin) |
| | | 56 | fc (PDGFRalin) |
| | | 56 | fc (PDGFRalin) |
| | | 64 | fc (PDGFRalin) |

| | | | |
|---|---|---|---|
| Figure 1K | Flow cytometry | 70 | fc (PDGFRa, MEFSK4) |
| | | 57 | fc (PDGFRa, MEFSK4) |
| | | 49 | fc (PDGFRa, MEFSK4) |
| | | 65 | fc (PDGFRa, MEFSK4) |
| | | 58 | fc (PDGFRa, MEFSK4) |
| | | 75 | fc (PDGFRa, MEFSK4) |
| Figure 2I | Hydroxyproline assay | 91 | kq (PDGFRa) |
| | | 98 | kq (PDGFRa) |
| | | 60 | kq (PDGFRa) |
| | | 96 | kq (PDGFRa) |
| | | 83 | kq (PDGFRa) |
| | | 79 | kq (PDGFRa) |
| | | 81 | kq (PDGFRa) |
| Figure 2J | SEM | 76 | kq (PDGFRa) |
| | | 83 | kq (PDGFRa) |
| | | 90 | kq (PDGFRa) |
| Figure 2L | Western blot (7 months post-induction) | 57 | wb (PDGFRa) |
| | | 72 | wb (PDGFRa) |
| | | 76 | wb (PDGFRa) |
| Figure 3 | Shotgun proteomics | 75 | kq (PDGFRa) |
| | | 74 | kq (PDGFRa) |
| | | 75 | kq (PDGFRa) |
| | | ~~33~~ | kq (PDGFRa) |
| Figure 4 | 10 week MI | 86 | kq (PDGFRa) |
| | | 72 | kq (PDGFRa) |
| | | 80 | kq (PDGFRa) |
| | | 82 | kq (PDGFRa) |
| | | 65 | kq (PDGFRa) |
| Figure 5 | 28d AngII/PE histology | 69 | kq (PDGFRa) |
| | | 74 | kq (PDGFRa) |
| | | 68 | kq (PDGFRa) |
| | | all samples 70-80 | tf |
| Figure 6 | 14d AngII/PE microarray | 55 | hq (PDGFRa) |
| | | ~~12~~ | hq (PDGFRa) |
| | | 56 | hq (PDGFRa) |
| | | 61 | hq (PDGFRa) |
| | | 66 | hq (PDGFRa) |
| | | 49 | hq (PDGFRa) |
| | PV loop analysis | 85 | kq (PDGFRa) |

| | | | |
|---|---|---|---|
| SFigure 1 G-H | | 47 | kq (PDGFRa) |
| | | 93 | kq (PDGFRa) |
| | | 93 | kq (PDGFRa) |
| | | 90 | kq (PDGFRa) |
| SFigure 3 | TAILS proteomics | 70 | kq (PDGFRa) |
| | | 88 | kq (PDGFRa) |
| | | 70 | kq (PDGFRa) |
| | | 88 | kq (PDGFRa) |
| | | ~~29~~ | kq (PDGFRa) |
| | | ~~56~~ | kq (PDGFRa) |

Abbreviations: tf, tomato fluorescence; gf, gfp fluorescence; wb, western blot; hq, qPCR from heart; kq, qPCR from kidney; fc, flow cytometry; strikethrough indicates excluded sample from study. Note: Perfusion techniques used for primary cardiomyocyte isolation resulted in poor RNA quality, thus no deletion efficiency could be evaluated for these samples.

**Supplemental Table 4. Cell-specific reagents for tissue/cell staining.**

| Antibody/reagent | Clone | Application (conc. ug/ml) | Vendor (cat no.) |
|---|---|---|---|
| Anti-collagen I | polyclonal | TS (10.0) | Genetex (GTX20292) |
| Anti-CD11b | M1/70 | FC (0.4) | BioLegend (101225) |
| Anti-CD140a | REA637 | FC (1.5) | Miltenyi (130-109-736) |
| Anti-CD31 | REA784 | FC (3.0) | Miltenyi (130-111-356) |
| Anti-CD45 | 30-F11 | FC (1.0) | BioLegend (103126) |
| Anti-collagen IV | polyclonal | TS (1.0) | Millipore (AB748) |
| Anti-collagen IV | polyclonal | WB (1.0) | Thermo Fisher (PA5-104508) |
| Anti-collagen VI alpha 1 | SD83-03 | WB (1.0) | Novus biologicals (NBP2-67825) |
| Anti-GAPDH | 6C5 | WB (3.0) | Thermo Fisher (AM4300) |
| Anti-Gr-1 | RB6-8C5 | FC (0.2) | BioLegend (108411) |
| Anti-laminin | polyclonal | TS (5.0) | Sigma (L9393) |
| Anti-laminin | polyclonal | WB (1.0) | Thermo Fisher (PA1-16730) |
| Anti-Ly6G | 1A8-Ly6g | FC (0.2) | eBioscience (12-9668-80) |
| Anti-mEF-SK4 | mEF-SK4 | FC (7.5) | Miltenyi (130-120-166) |
| Anti-PDGFRα | polyclonal | TS (2.0) | R&D systems (AF1062) |
| Anti-PDGFRα | D1E1E | WB (0.023) | Cell signaling (3174) |
| Anti-vimentin | polyclonal | TS (10.0) | Abcam (ab45939) |
| Wheat germ agglutinin | NA | TS (5.0) | Vector Labs (B-1025) |
| TS: tissue staining; WB: Western blot; FC: Flow cytometry | | | |

**Supplemental Table 5. Primers used for qRT-PCR.**

| Gene | Forward sequence (5'-3') | Reverse sequence (5'-3') |
|---|---|---|
| *Pdgfra* | GTCGTTGACCTGCAGTGGA | CCAGCATGGTGATACCTTTGT |
| *Tcf21* | GGCCAACGACAAGTACGAGA | GCTGTAGTTCCACACAAGCG |
| *Vim* | CCAACCTTTTCTTCCCTGAA | TGAGTGGGTGTCAACCAGAG |
| *Col1a1* | AATGGCACGGCTGTGTGCGA | AACGGGTCCCCTTGGGCCTT |
| *Col1a2* | GGCCCCCTGGTATGACTGGCT | CGCCACGGGGACCACGAATC |
| *Dpep1* | CCATCTGTGCTTCGGACTCAT | CCAAGGCAAGTCGTTGTGC |
| *Dkk3* | CTCGGGGGTATTTTGCTGTGT | TCCTCCTGAGGGTAGTTGAGA |
| *Col5a1* | TGAGTCTGGTTTTCCCGAGGA | GCCCTGCTCATTGTAAATGGAGA |
| *Mdk* | TGGAGCCGACTGCAAATACAA | GGCTTAGTCACGCGGATGG |
| *Pdgfrl* | CGGACTTCTGTTGCTACACGA | TGGTTGGTTTGATCCTGTTTTCT |
| *18s* | GTAACCCGTTGAACCCCATT | CCATCCAATCGGTAGTAGCG |
